# Quadratic and adaptive computations yield an efficient representation of song in *Drosophila* auditory receptor neurons

**DOI:** 10.1101/2021.05.26.445391

**Authors:** Jan Clemens, Mala Murthy

## Abstract

Sensory neurons transform stimuli through multiple adaptive and nonlinear computations. In *Drosophila*, auditory receptor neurons adapt to the mean and intensity of acoustic stimuli, shift their frequency tuning with sound intensity, and employ a quadratic nonlinearity. Although each of these computations is thought to improve sensory encoding, their combination could produce a highly ambiguous neural code from which behaviorally relevant information is difficult to extract. Here, we combine electrophysiology and computational modeling to determine how these computations jointly encode natural acoustic signals such as courtship song.

We show that a model consisting of a quadratic filter followed by divisive normalization accurately reproduces population responses to both artificial and natural sounds. The model further localizes adaptive temporal filtering to antennal mechanics and variance adaptation to divisive normalization downstream of the receptor mechanics. For arbitrary broadband stimuli, these computations generate an ambiguous representation from which carrier frequency and amplitude cannot be reliably recovered. In contrast, the representation of courtship song is simple and robust. Quadratic filtering enhances encoding of the song envelope while preserving a straightforward temporal code for the carrier through frequency doubling. Divisive normalization removes slow intensity fluctuations arising during social interactions while preserving the rapid envelope fluctuations that define song structure.

Together, our results show that adaptive and nonlinear computations that appear to complicate sensory coding for arbitrary stimuli instead generate a robust and easily decoded representation of behaviorally relevant signals by exploiting their natural statistics.

**Author Summary:** Animals do not need to encode every aspect of the physical world equally well. Instead, sensory systems must preserve the features of signals that matter for behavior. We studied the auditory receptor neurons of fruit flies, which respond to courtship song and adapt when sound intensity changes. We developed a model that captures several computations performed by these neurons and used it to ask how they affect the representation of natural song. The model shows that changes in the mechanics of the fly’s antenna adjust frequency tuning, whereas a later normalization step reduces the effects of slow changes in sound intensity. A quadratic transformation doubles sound frequencies. Although this can make arbitrary sounds ambiguous, it preserves the carrier frequency of fly song and improves representation of its envelope, the slower pattern of sound intensity. Together, these computations make the neural representation of courtship song robust to noise and intensity changes while retaining the information needed to recognize song patterns.

## Introduction

Sensory receptor neurons transform physical stimuli into neuronal representations. Constraints on neuronal dynamic range and bandwidth result in neural codes that represent some aspects of the stimulus at the cost of others, typically through multiple nonlinear and adaptive computations. For instance, sensory neurons are known to adapt to the statistics of their inputs: Visual systems adapt to the luminance and contrast of a visual scene, and auditory systems to the frequency and intensity statistics of the soundscape [1, 2]. Adaptation creates energy-efficient representations and improves stimulus discrimination [3–7]. Adaptation can act on two properties of the mapping from stimulus to neural response: 1) how inputs are integrated in time and/or space (filtering), and 2) how the integrated input is transformed into spikes (gain control) [7–13]. However, while adaptive filtering and gain control improve information transmission, they also remove information, for instance about absolute light or sound intensity levels. This introduces ambiguity for neural computations in downstream circuits since the meaning of a spike—the pattern and magnitude of the stimulus it represents—will depend on the stimulus history [4, 14–17]. This ambiguity is desirable when it introduces invariances to intensity or contrast, but it may distort neural readouts if the adaptation affects behaviorally relevant stimulus features [4, 18]. Whether adaptation is beneficial therefore depends not only on information transmission, but also on which stimulus features downstream circuits must decode.

Like adaptation, other nonlinear transformations are beneficial for making explicit some stimulus features while sacrificing others. For instance, the recognition of acoustic signals—such as speech—typically relies on two features of a sound: 1) the carrier, or fast oscillations that define the fine structure of the waveform, and 2) the envelope, relatively slower modulations of the intensity or variance of the carrier. A quadratic output nonlinearity produces responses that depend on the magnitude but not on the sign of an input stimulus. This operation is considered crucial for extracting the envelope, but it introduces ambiguity about the structure of the carrier. Importantly, the impact of this ambiguity on decoding depends on the stimulus: For spectrally simple, pure-tone carriers, a quadratic nonlinearity produces frequency doubling responses, from which the stimulus frequency can be easily recovered. However, for spectrally complex carriers, the frequency content of the stimulus cannot generally be recovered after a quadratic nonlinearity. Thus, evaluating sensory codes using generic stimulus ensembles can be misleading. The consequences of adaptation and nonlinear transformations depend on the statistics of the stimuli being encoded and on the stimulus features that must ultimately drive behavior [19]. The same computation may therefore simplify neural readout for natural signals while introducing ambiguity for arbitrary stimuli.

We address this issue in the auditory receptor neurons of *Drosophila melanogaster*. Hearing plays a central role in *Drosophila* courtship [20]: Males chase females and produce a courtship song by vibrating their wing. The male song contains two major modes: The sine song consists of sustained oscillations with a frequency around 150 Hz, and the pulse song consists of regular trains of short pulses that can have two distinct shapes [21, 22]. Females, and nearby males, use both the spectral and the temporal properties of the song to guide social interactions [23–26]. In addition to acoustic communication, flies use acoustic cues for threat detection based on sudden increases in the sound envelope [27]. Mechanoreceptor neurons in the fly’s antenna—its “ear”—must therefore preserve information about the carrier and the envelope of the song for further processing downstream [28–33]. Flies detect sound using the arista, a feathery extension of the antenna (Fig. 1A). Sound-induced antennal vibrations, but also slow antennal movement induced by wind and gravity, activate a diverse population of stretch-sensitive mechanoreceptors in the antenna’s second segment—the Johnston’s organ neurons (JONs) [34–36]. While some types of JONs encode slow antennal movement induced by wind and gravity (JONs C and E) [36], the JON-A and JON-B subpopulations respond best to fast, sound-induced antennal vibrations in the frequency range of the courtship song (100–400 Hz) and act as the auditory receptor neurons. These auditory JONs perform several adaptive and nonlinear computations that are thought to improve sound encoding (Fig. 1B): JONs adapt to the mean and to the variance of receiver movement [27, 37, 38]. Mean adaptation renders responses invariant to slow antennal movement induced by wind and gravity. Variance adaptation compensates for slow intensity fluctuations, which arise during the dynamic social interactions caused by the directional and distance-dependent mechanics of sound reception [39, 40]. The extracellularly recorded bulk spiking activity of JONs A and B—the compound action potential (CAP) [27, 34, 38] (Fig. 1A)—exhibits frequency doubling responses for sinusoidal stimuli. For instance, a 300 Hz sinusoidal stimulus evokes 600 Hz oscillations in the CAP [27, 41, 42] (Fig. 1B). This frequency doubling can be reproduced mathematically by squaring the sinusoidal stimulus, indicating that sound undergoes a quadratic transformation across the JON population. JONs also exhibit adaptive temporal filtering: the cutoff frequency of the antennal receiver increases with intensity [43] and this effect is linked to the mechanical amplification of low-amplitude sounds in JONs [44–47]. How these computations interact to encode natural sounds remains unknown. Addressing this question requires a computational model that simultaneously captures mean and variance adaptation, quadratic nonlinearity, and adaptive temporal filtering while reproducing responses to arbitrary dynamic stimuli.

**Figure 1:**
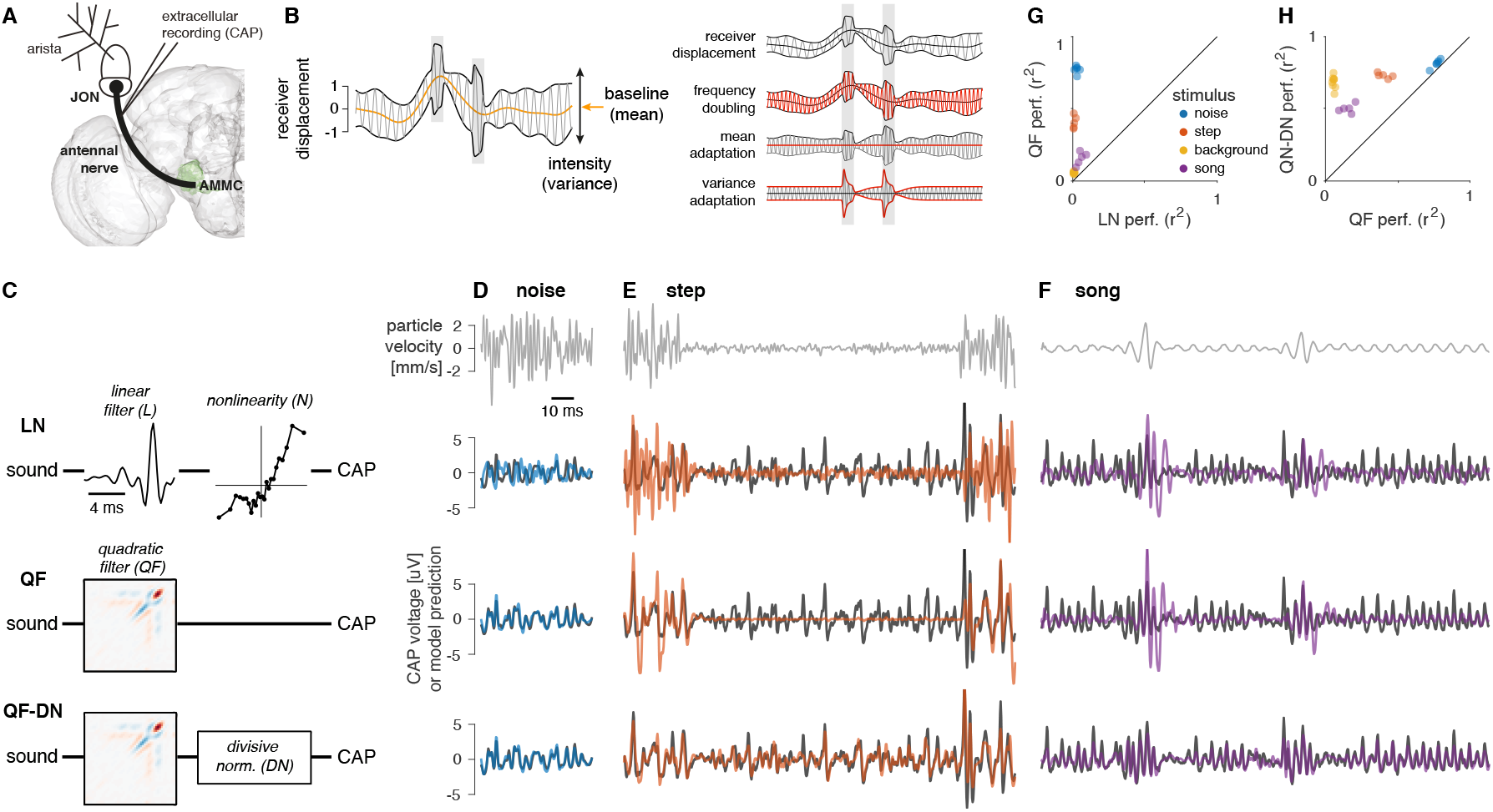
A quadratic filter with a divisive normalization stage reproduces JON responses to a wide range of stimuli. **A** Schematic of the early auditory system of Drosophila and recording location. Sound-induced vibrations of the arista induce antennal rotations and activate antennal mechanosensory neurons—the Johnston’s organ neurons (JONs). The compound action potential (CAP) represents the activity of sound-responsive JONs and can be recorded using an electrode inserted into the joint between the first and second antennal segments. **B** Left: Fast oscillation (carrier, gray line) with a slowly varying baseline (mean, orange line) and slow and fast (gray boxes) fluctuations of the intensity (variance, black line). Right: Schematic illustration of four JON computations evident from the CAP (the response feature that changes in subsequent rows is marked in red). **C** Structures of the tested models. The linear-nonlinear (LN, top) model consists of a linear filter followed by a static nonlinearity. The quadratic filter (QF, middle) model consists of a quadratic filter without a nonlinearity. The QF-DN (bottom) model is a quadratic filter with a divisive normalization (DN) stage. Filters and nonlinearities represent examples of models fitted to CAP recordings. **D, E, F** CAP (black) and model (colored lines) responses of the three different models in C (rows 2–4) for different stimuli (top row). “Noise” (D) corresponds to band-pass-filtered white noise with constant intensity. “Step” (E) is noise with stepwise changes in intensity every 100 ms. “Song” (F) corresponds to a male courtship song and contains sustained oscillations, called sine song, and trains of transient pulses. Data from one representative fly. **G, H** Comparison of the performance (coefficient of determination *r* ^*2*^ between the CAP and model prediction) for the different models and stimuli in C–F (color coded, see legend). Points correspond to recordings from individual flies. The LN model performs poorly for all stimuli. The QF model performs well for stimuli with a constant intensity (noise), but poorly for stimuli with dynamic intensity profiles (step, background, song). The QF-DN model performs well for all stimuli. “Background” is not shown in D–F and is a noise stimulus of mostly constant amplitude with brief changes in intensity (see Fig. S6). Numbers of flies for noise/step/background/song are 6/5/9/5.

However, existing JON models focus on individual computations, such as active mechanical amplification [27, 48] or the interplay between mean and variance adaptation [38]. We therefore developed and fitted a comprehensive computational model of the JONs based on CAP responses to artificial and natural stimuli. We based the model on CAP recordings because they currently provide the only population readout of JON activity with sufficient temporal resolution to capture all of these computations. The model reproduces all major features of the responses for a wide range of stimuli including the courtship song and generates hypotheses about the mechanisms that implement the individual computations and about the contribution of individual processing steps to the representation of sound in JONs. Specifically, by manipulating individual components of the model, we identify the contribution of each computation to sound encoding. We find that the combination of adaptive and nonlinear transformations produces a representation of courtship song that is both robust and easy to decode, allowing behaviorally relevant features of the carrier and envelope to be recovered with little ambiguity.

## Results

### A model with a quadratic filter and divisive normalization reproduces CAP responses

To fit a model that captures the multiple computations performed by JONs, we recorded CAP responses to stimuli with diverse carrier and envelope dynamics (Fig. 1C): Gaussian noise band-pass filtered between 100 and 1000 Hz (hereafter termed “noise”) with either constant intensity (Fig. 1D) or step changes in amplitude (Fig. 1E); and the natural courtship song, which exhibits narrow-band carriers and strong amplitude modulations (Fig. 1F).

A common framework for describing the stimulus-response mapping of sensory neurons is the linear-nonlinear (LN) model [49, 50] in which the stimulus is 1) linearly filtered to account for temporal integration properties and 2) transformed by a static nonlinearity to account for neuronal threshold or saturation (Fig. 1C, top). However, the LN model fails to reproduce the nonlinear CAP responses (Fig. 1D–F, top; Fig. 1G). Replacing the linear filter with a quadratic filter (QF, Fig. 1C, middle) accurately captures the fine structure of responses to constant-intensity noise but fails for stimuli with dynamic intensity changes (Fig. 1D–G). For instance, the QF does not reproduce the response transients following intensity steps or the relative intensity invariance of the steady-state responses (Fig. 1E, F).

To account for this adaptive gain control, we added a divisive normalization (DN) stage after the QF (Fig. 1C, bottom). The DN stage was placed downstream of the QF because variance adaptation arises in the JONs after mean adaptation and frequency doubling [27, 38]. Specifically, the DN stage low-pass filters the rectified QF output to generate an adaptation signal, which divisively scales the QF output. Adding the DN stage substantially improved model performance for stimuli with dynamic envelopes, particularly courtship song (Fig. 1E, F, H).

A comparison of model performance using the Bayesian Information Criterion, which penalizes complex models by considering the number of model parameters, confirms that a model with a quadratic filter and divisive normalization reproduces JON responses best (Fig. S1E). We refer to this two-stage model as QF-DN. Because the model accurately reproduces responses to a wide range of stimuli, it provides a framework for dissecting how quadratic filtering and divisive normalization contribute to the representation of courtship song. We begin by examining how the quadratic filter shapes the representation of the song’s carrier and envelope.

### The quadratic filter is a frequency-dependent encoder of sound carrier and envelope

A linear filter corresponds to a set of weights, *h*(*τ*), applied to the stimulus *s* (*t − τ*) at different delays *τ* : *r* (*t*) = ∑_*τ*_ *h*(*τ*)*s*(*t − τ*) (Fig. 1C, top). A quadratic filter also constitutes a set of weights, *H*_*2*_ (*τ*_*1*_, *τ*_*2*_), but these act on products of stimulus values at different delays *τ*_*1*_ and 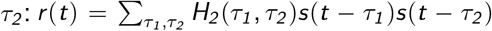 (Fig. 2A). While a linear filter can often be inferred directly from its set of weights—for instance, purely positive filters smooth the stimulus while filters with positive and negative lobes are band-pass filters—the stimulus selectivity of a quadratic filter is much less intuitive. To obtain an interpretable representation, we transformed the QF into a bank of linear filters with quadratic output nonlinearities using an eigenvalue decomposition of its weight matrix *H*_*2*_ (Fig. 2B, S1) [51, 52]. The outputs of the individual filters (Fig. 2C, D) are then linearly combined to predict the CAP: *r* (*t*) = ∑_*i*_ *σ*_*i*_ [∑_*τ*_ *s*(*t − τ*)*v*_*i*_ (*τ*)]^2^. The eigenvectors, *v*_*i*_, define the linear filters, whereas the eigenvalues, *σ*_*i*_, determine their contribution to the overall response (Fig. 2C, E). The full model also contains linear and constant terms. However, these contributed negligibly to model performance (Fig. S1D) and were omitted from the analyses below. This decomposition allows the quadratic filter to be interpreted as a set of frequency-selective channels whose outputs are rectified and summed.

**Figure 2:**
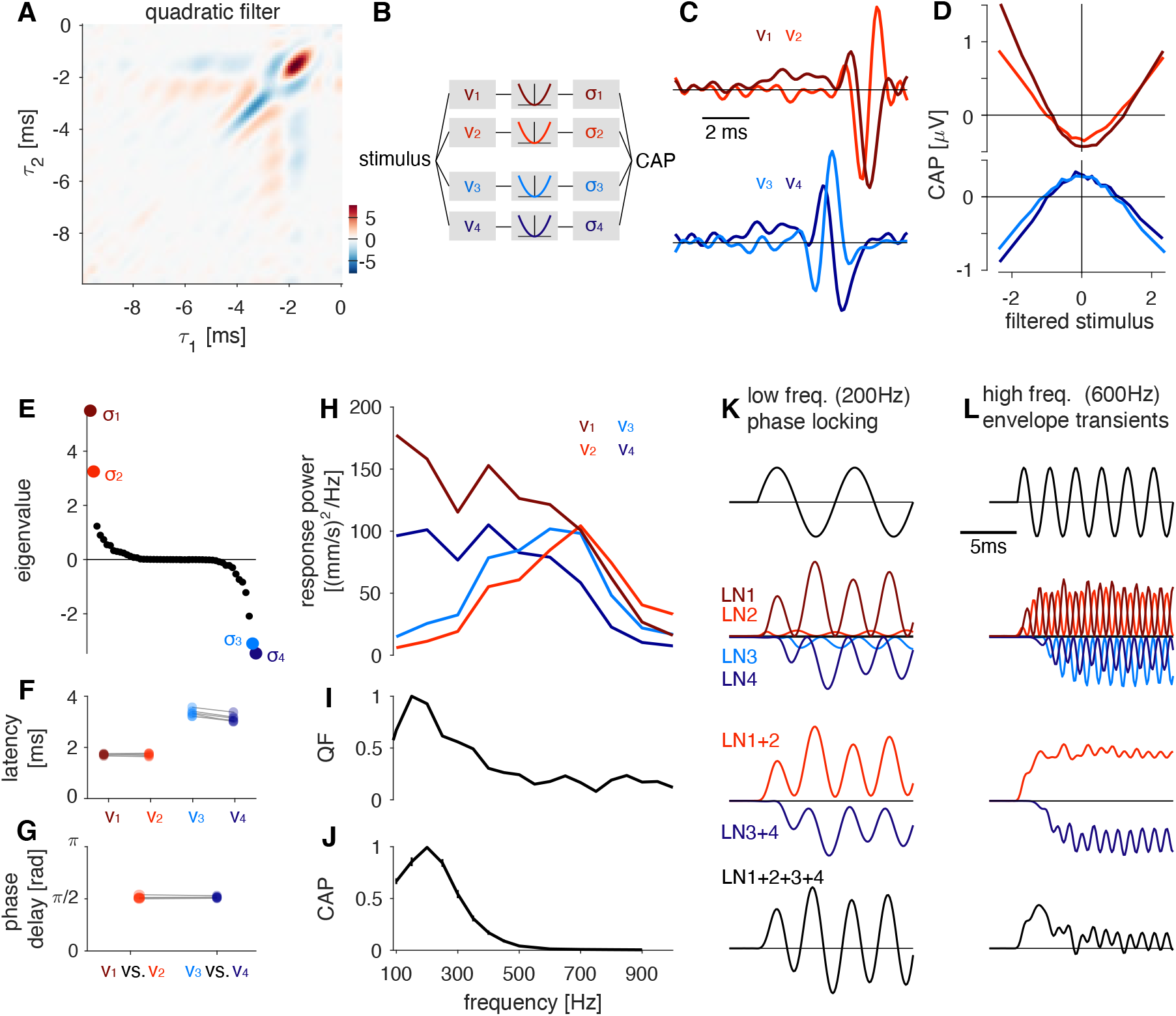
The quadratic filter differentially encodes low and high frequencies. **A** The quadratic filter (QF) is a matrix *H*_*2*_ (*τ*_*1*_, *τ*_*2*_) that contains weights for the product of stimulus values for different *τ*. *τ* = *0* corresponds to the time of the response and negative values to the time before the response. The filter was fitted to CAP responses to noise presented at 0.5 mm/s. Filter matrix values are color coded (see color bar). **B** The QF can be transformed into a bank of linear filters with quadratic nonlinearities (LN models, pictograms) using eigenvalue decomposition (Fig. S1). Eigenvectors *v*_*i*_ correspond to the filters (see C) and the eigenvalues *σ*_*i*_ correspond to the weights for each eigenvector (see E) in the filter bank. Together with their associated nonlinearities and eigenvalues, the four eigenvectors with the largest-magnitude eigenvalues form four LN models (LN1–4) that reproduce the performance of the full quadratic filter (Fig. S1D). **C** Eigenvectors associated with the largest positive (top, *v*_*1*_ and *v*_*2*_, red hues) and negative (bottom, *v*_*3*_ and *v*_*4*_, blue hues) eigenvalues (see E). Each vector was normalized to unit norm. **D** The relationship between the stimulus filtered by the eigenvectors in C and the response is quadratic. *v*_*1*_ and *v*_*2*_ form an “excitatory” filter pair since filter outputs with large magnitude increase the CAP. By contrast, *v*_*3*_ and *v*_*4*_ form a “suppressive” filter pair. Curves correspond to the average CAP for binned values of the filtered stimulus. **E** Sorted eigenvalues of *H*_*2*_. Values for *σ*_*1*_ –*σ*_*4*_ are highlighted in color. **F, G** Latency of the filters (F) and phase delays between the filters in a pair (G). Dots correspond to individual flies; lines connect data from the same individual. N=6 flies. See also Fig. S2B–D. **H** Frequency transfer functions of each filter in C. Each filter pair contains a filter that responds to low frequencies (*v*_*1*_, *v*_*4*_) and a filter that prefers high frequencies (*v*_*2*_, *v*_*3*_). **I, J** Frequency tuning of the phase-locked component at twice the stimulus frequency in the model (I) and the CAP (J). Tuning curves were normalized to peak at 1.0. **K, L** Responses of the 4 individual LN (2nd row), filter pairs (3rd row), and the full QF (4th row) to 200 Hz (K) and 600 Hz (L) sinusoidal stimuli (1st row). For low frequencies (K) the four filters combine to produce sustained oscillations at twice the stimulus frequency. For high frequencies (L) the QF phase-locks weakly and responds strongly to the stimulus onset. All panels except F, G show data from one representative fly. Filter structure is very similar across individuals (see F, G).

This eigenvalue decomposition provides a compact representation of the computation implemented in JONs: Only four filters corresponding to the four largest-magnitude eigenvalues are sufficient to reproduce the performance of the QF (Fig. S1B–D). The four filters are each well approximated by Gabor filters (Fig. S2A–B). Because these filters contain both positive and negative lobes, they respond only weakly to static stimuli and are tuned to fluctuating inputs (Fig. 2C, S2E). This band-pass structure naturally accounts for mean adaptation in JONs, which suppresses responses to static antennal deflections. Beyond this common structure, the four filters are organized into two filter pairs: an excitatory pair with short latency and a suppressive pair with longer latency (Fig. 2C–F). The suppressive pair resembles a delayed, sign-inverted version of the excitatory pair (Fig. 2C). Within each pair, the two filters are phase-shifted by 90° and thus form a quadrature pair (Fig. 2G). Importantly, this filter organization emerges regardless of the stimulus ensemble used for model fitting (Fig. S3A–D).

Quadrature-pair filters are well known from auditory nerve fibers in vertebrates [52], complex cells in the primary auditory and visual cortex of vertebrates [53, 54], and motion-sensitive cells in the vertebrate visual cortex or the *Drosophila* optic lobe [55]. Canonical quadrature pairs extract the stimulus envelope because the outputs of the two filters combine to a phase-invariant, smoothly varying signal proportional to the stimulus energy [52, 56]. At first glance, this interpretation appears inconsistent with JON responses, which phase-lock to sounds (Fig. 1D–F). This apparent contradiction is explained by an asymmetry in the frequency tuning of the filters within each pair (Fig. 2H). Quadrature pairs act as envelope detectors only for carrier frequencies at which both filters respond strongly. In JONs, this occurs only above approximately 400 Hz (Fig. 2H, L). In this high frequency range, each quadrature pair encodes the stimulus envelope. Because the suppressive pair is delayed relative to the excitatory pair (Fig. 2C, F), the combined response emphasizes transient increases in stimulus energy (Fig. 2L). Thus, the filter structure accounts for key aspects of variance adaptation, including the transient responses at high frequencies [57–59]. Below 400 Hz, by contrast, only one filter in each quadrature pair responds strongly (Fig. 2H, K). Consequently, the QF effectively behaves as two linear filters followed by quadratic nonlinearities. In this regime, it produces sustained responses that phase-lock at twice the carrier frequency, matching the behavior of the CAP (Fig. 2I–K). Thus, the QF reveals two distinct coding regimes in JONs: above 400 Hz it extracts changes in stimulus envelope, whereas below 400 Hz it preserves information about the sound carrier through sustained, phase-locked frequency-doubling responses [42].

Overall, the quadratic filter accounts for several hallmark computations of JONs, including frequency doubling, mean adaptation, and variance adaptation at high frequencies. We next asked whether the filter itself adapts by examining how its structure changes with sound intensity.

### Adaptive temporal filtering arises from adaptive antennal mechanics

To characterize adaptive processes that shape filter properties, we analyzed QFs fitted to CAP responses to noise stimuli over a wide range of sound intensities (1/16 mm/s to 4 mm/s, Fig. 3A, B, S4A, B). Eigenvalue decomposition of these QFs revealed only modest changes in the relative timing and weight (eigenvalues) of the Gabor filters (Fig. 3C, S4A). The dominant effect, however, was a narrowing of the negative lobe of all four Gabor filters, increasing their cutoff frequencies by approximately 200–500 Hz across the tested intensity range (Fig. 3D). Thus, the temporal filtering implemented by the QF becomes progressively faster with increasing sound intensity, consistent with predictions from optimal coding theory [9, 10] and observations in other systems [2, 13].

**Figure 3:**
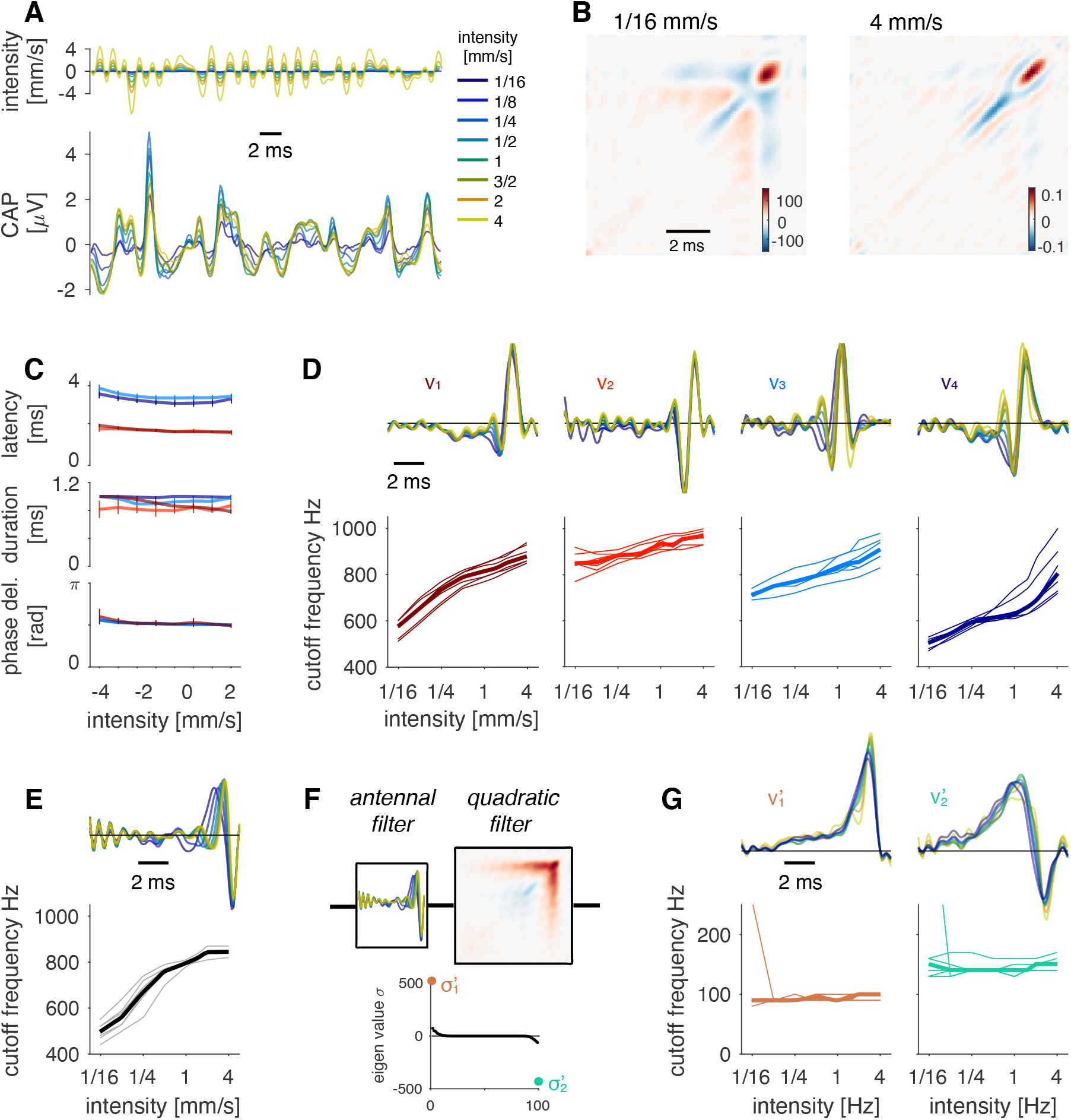
Antennal mechanics produce adaptive filtering in JONs. **A** CAPs (bottom) for noise patterns (top) of different intensities (color coded, see legend). **B** Structure of the quadratic filter (see color bar) at the lowest (left) and the highest (right) intensity tested. The filter structure along the main diagonal narrows with intensity. **C** Latencies (top), durations (middle), and phase delays (bottom) of the four leading eigenvectors (filters) *v*_*1*_ –*v*_*4*_ (color coded as in D) change little with intensity. Lines and error bars represent median ± std over 6 flies. **D** Shape (top, intensity color-coded, see legend in A) and cutoff frequency (bottom) of the filters *v*_*1*_ –*v*_*4*_ for different inten-sities. Filters (top) are from one representative fly. Cutoff frequencies for all filters increase with intensity (*r > 0* .*95, p < 4* × *10* ^*−4*^ for all *v*). **E** Shape (top, example from one fly, intensity color-coded, see legend in A) and cutoff frequency (bottom) for the antennal filter for different intensities. The filter describes the transformation from sound stimulus to antennal movement, and is measured using laser Doppler vibrometry. The cutoff frequency increases with intensity (*r* = *0* .*97, p* = *7* × *10* ^*−5*^). **F** Accounting for antennal mechanics greatly simplifies the filter structure. In the antQF model, the sound was first filtered with the intensity-dependent antennal filter (top left) before fitting the QF (top right). The eigenvalue spectrum (bottom) of the resulting quadratic filter reveals that two filters (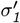 and 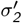) are sufficient to reproduce the structure of the antQF (cf. Fig. 2E). **G** Shape (top, example from one fly, intensity color-coded, see legend in A) and cutoff frequency (bottom) for the two dominant eigenvectors of the antQF, 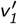 and 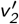. The filters are slower and change little with intensity compared to the filters of the original QF (compare D). r=0.18, p=0.7 (left), r=0.81, p=0.016 (right), slope is negligible in both cases (<1.7 Hz/intensity doubling). The outliers stem from weak CAP responses to the lowest intensity in one fly (Fig. S4B). N=6 flies for C–G. P-values from linear fits to the mean across flies. Thin lines in D, E, G (bottom) correspond to individual flies, the thick line to the median across 6 flies.

What gives rise to this adaptive temporal filtering? Previous studies showed that the frequency tuning of the antennal receiver depends on sound intensity because of active processes driven by mechanotransducer gating in JONs [43, 47, 60, 61]. Using laser Doppler vibrometry, we confirmed that the antenna’s cutoff frequency increases with sound intensity (Fig. 3E, S4C). We next asked whether these adaptive antennal mechanics fully account for the observed changes in the QF. To this end, we decomposed the filter into a mechanical and a neuronal stage: We first passed the stimulus through the intensity-dependent antennal filter measured by vibrometry (Fig. 3E), thereby accounting for the mechanical transformation of sound. We then fitted a new quadratic filter to this mechanically filtered stimulus (Fig. 3F), such that the new filter captures only the remaining neuronal computations.

We refer to this decomposition as the antQF (antennal QF) model. Accounting for intensity-dependent antennal filtering dramatically simplified the quadratic filter (Fig. 3F): The resulting filter retained only the slow components associated with the timing of the excitatory and suppressive filter pairs present in the original model (Fig. 3B). Eigenvalue decomposition showed that the antQF is well described by only two eigenvectors, rather than four. Moreover, these eigenvectors lack the fast oscillatory structure present in the original QF (Fig. 3G, compare Fig. 3D), indicating that this structure is fully captured by the mechanical antennal filter. Unlike the original QF, both the shape and frequency tuning of the antQF are intensity invariant.

These results demonstrate that adaptive temporal filtering in JONs arises from adaptive antennal mechanics driven by mechanotransducer gating. The antQF simply decomposes the original QF into an intensity-dependent linear mechanical stage followed by an intensity-invariant quadratic neuronal stage (Fig. 3F). Although the two models are computationally equivalent and produce identical outputs, this decomposition shows that the adaptive frequency tuning previously observed in JONs is inherited from antennal mechanics rather than arising from changes in neuronal filtering.

### Divisive normalization reproduces the strength and dynamics of variance adaptation

Antennal filtering reproduces the adaptive temporal filtering of JONs, but not the variance adaptation observed in the CAP (Fig. S5). First, whereas CAP responses remain nearly invariant over a wide range of sound intensities, antennal responses scale almost linearly with the stimulus intensity. Second, CAP responses exhibit prominent transients following intensity steps that are absent from antennal responses. Although the QF captures some aspects of variance adaptation at high frequencies (Fig. 2L), it fails to account for adaptation across the full frequency range (Fig. 2K), including responses to courtship song (Fig. 1F, H). Thus, as indicated by our model comparisons (Fig. 1C–H), an additional divisive normalization (DN) stage is required to reproduce the adaptive gain of JONs [11]. In the DN stage (Fig. 4A), the output of the quadratic filter is divided by an adaptation signal—a running estimate of the response gain—obtained by rectifying and low-pass filtering the QF output. The DN stage has two free parameters that determine the strength (*θ*) and speed (*τ*) of adaptation. Note that *θ* is a phenomenological parameter that captures the effective adaptation strength for each stimulus class, not the underlying biophysical parameter.

**Figure 4:**
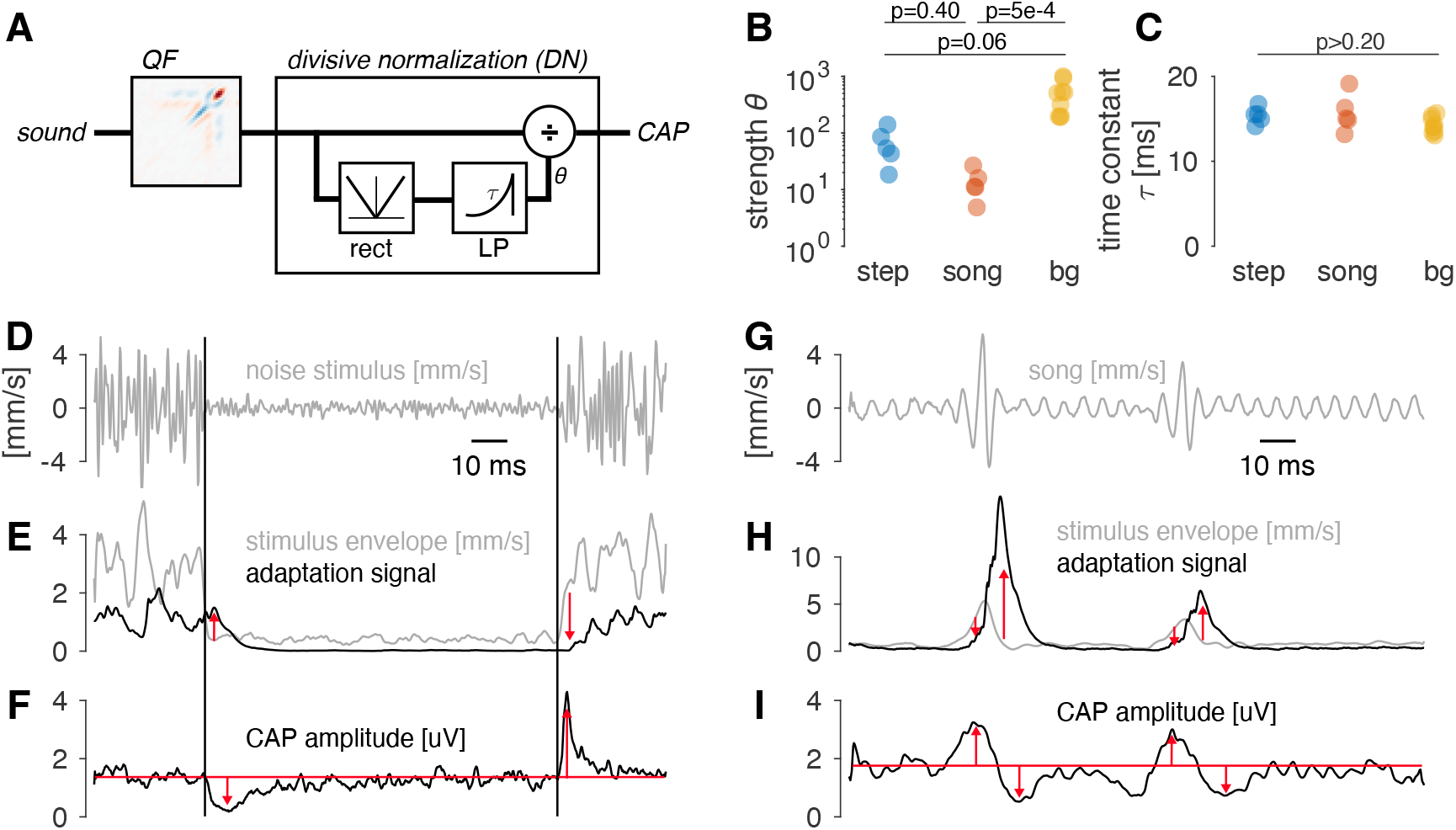
Divisive normalization explains variance adaptation and the transient responses to changes in intensity. **A** Internal structure of the divisive normalization stage used to model variance adaptation. The output of the QF is rectified and low-pass filtered to obtain an adaptation signal which divides the output of the QF. *τ* and *θ* control adaptation speed and strength, respectively. **B, C** Parameters of the model fitted to responses to different stimulus types (“bg” = background stimulus type). Adaptation strength *θ* (B) changes with the stimulus type but is ≫ *1* .*0* for all stimuli, which indicates near complete adaptation for all types. Adaptation time constants *τ* (C) do not change with the stimulus type. Dots correspond to individual flies. The p-values are from Tukey-Kramer post-hoc tests after a Kruskal-Wallis test. N=5/9/5 flies for step/background/song (different populations of flies from Fig. 3). **D, E, F** Stimulus (D), stimulus envelope (E, grey) and adaptation signal from the model (E, black). The adaptation signal lags the changes in stimulus intensity (red arrows). This induces transients in the CAP (F, black) due to over- and un-dercompensation of the stimulus intensity (red arrows). Vertical black lines in D–F indicate the switches in intensity. The horizontal red line in F depicts the CAP’s steady-state response. **G, H, I** Same as D–F but for song. See also Fig. S5.

Adaptation is stronger for artificial stimuli than for courtship song because artificial stimuli remain at constant intensity for long periods, allowing adaptation to reach steady state (Fig. 4B, S6F, G). By contrast, the rapidly fluctuating envelope of courtship song prevents adaptation from fully developing, leading to a lower effective adaptation strength.

The adaptation time constant *τ* is approximately 15 ms for all stimulus classes (Fig. 4C). This timescale reproduces the transient CAP responses for stimuli with dynamic envelopes (Fig. 4D–I). Following abrupt intensity changes, the adaptation signal lags behind the stimulus because it is generated by low-pass filtering the QF output (Fig. 4E, H). Consequently, gain compensation transiently underestimates increases and overestimates decreases in sound intensity (Fig. 4F, I), giving rise to the observed response transients. Notably, in the CAP, transients are slower for negative than for positive steps in intensity, and our adaptation model shows that these asymmetric dynamics arise from the multiplicative nature of adaptation with a single time constant parameter (Fig. S6A–E).

Overall, the QF-DN model accounts for the adaptive and nonlinear responses of JONs using just two computations: quadratic filtering and divisive normalization. Each computation improves one aspect of sensory coding while potentially degrading another. Quadratic filtering, which produces frequency doubling, distorts the representation of the sound carrier. Adaptive temporal filtering could further increase ambiguity because the transmission of different frequencies depends on stimulus intensity. Finally, variance adaptation may be too slow—or too fast—to efficiently encode transient communication signals such as pulse song. We therefore asked whether these apparent costs are offset by the statistics of natural courtship song.

### Quadratic filtering enhances the representation of the song envelope

We first asked whether the quadratic filter improves the representation of the song envelope. Canonical quadrature-pair filters extract the sound envelope [56]. However, the JON filter does not operate as a canonical envelope encoder over the low-frequency range occupied by courtship song (Fig. 2H–L). Instead, it behaves like a linear filter followed by a quadratic nonlinearity, producing frequency doubling responses in which each cycle of the carrier generates two response cycles (Fig. 1B, 2K). We hypothesized that this frequency doubling could nevertheless improve envelope coding by doubling the temporal sampling of the carrier and thereby better preserving rapid envelope fluctuations. To test this hypothesis, we set up two simple encoders: one that squares the stimulus just as in JONs and one that simply thresholds the song waveform (Fig. 5A).

**Figure 5:**
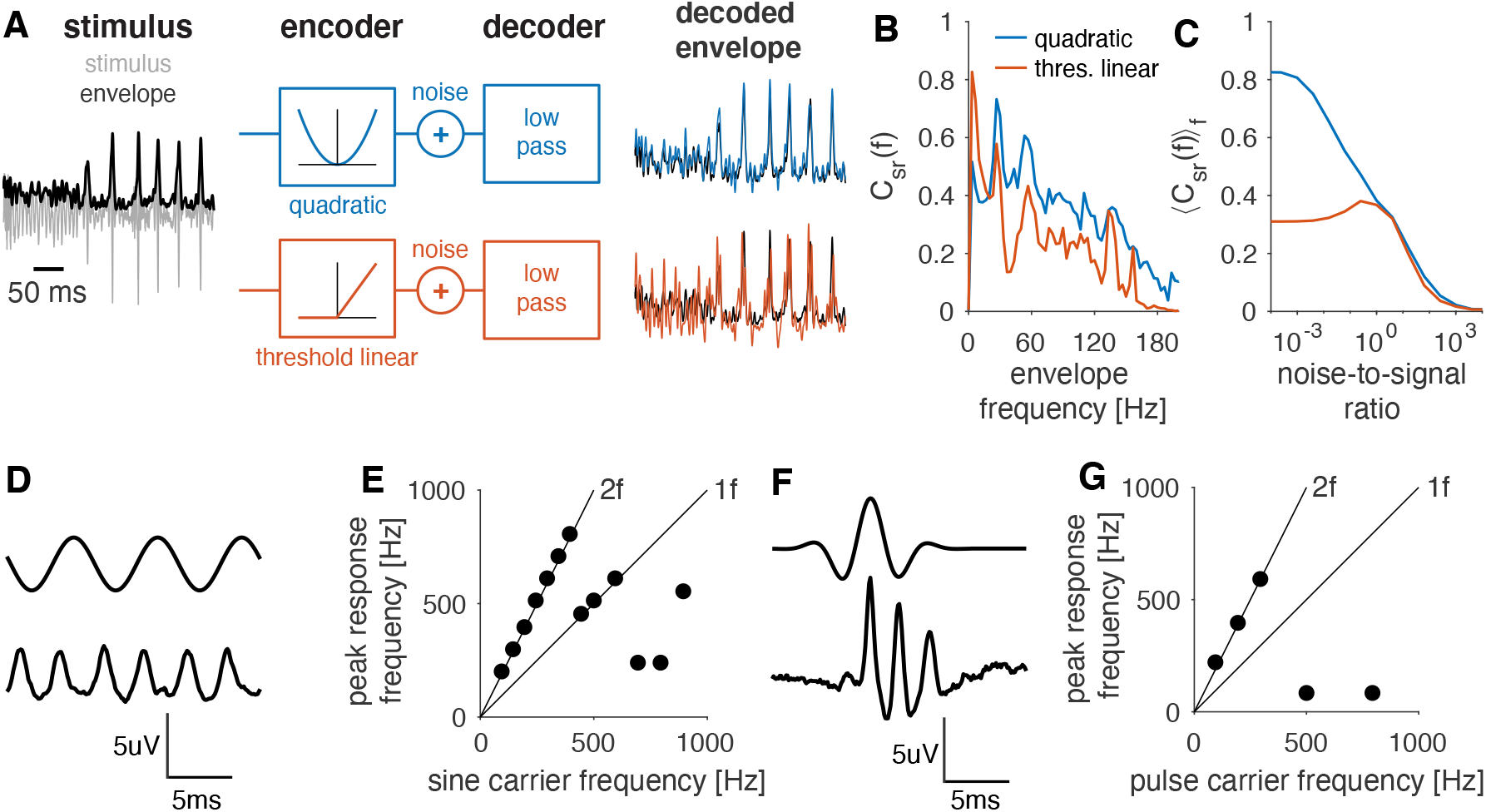
Quadratic filtering improves the representation of the sound envelope and yields a simple scheme for decoding the carrier frequency of song. **A** Encoder-decoder models used to test the role of a quadratic nonlinearity. The song (grey, envelope in black) is passed through a quadratic (blue, top) or a threshold-linear function (red, bottom). Then noise is added, and the response is low-pass filtered to reconstruct the stimulus envelope. Colored lines depict the envelope reconstruction from the quadratic (blue) and the threshold-linear (red) encoder, black lines show the envelope of the original stimulus. **B** Reconstruction success (coherence *C*_*sr*_ (*f*) between original and reconstructed envelope) for different frequency components of the envelope for quadratic (blue) and threshold-linear (red) encoders. The quadratic encoder wins for nearly all frequencies. The noise-to-signal ratio was 0.25. **C** Integral coherence over the range from 1–60 Hz for quadratic (blue) and threshold-linear (red) encoders for different noise-to-signal ratios. This frequency range includes the behaviorally relevant interval between song pulses (25–30 Hz). The quadratic encoder wins in particular in the low-noise regime. Results are similar when integrating the coherence over the full frequency range. **D** 150 Hz sine stimulus (intensity 2 mm/s) (top) and CAP (bottom). **E** Dominant frequency of the CAP response for sinusoidal stimuli with different frequencies. Lines correspond to peaks at the stimulus frequency (1f) and its double (2f) (see Fig. S7A). **F** Pulse with a carrier frequency of 200 Hz (intensity 2 mm/s) (top) and CAP (bottom). **G** Same as E but for pulses with different carrier frequencies (see Fig. S7B).

We then quantified how well an optimal linear decoder could reconstruct the song envelope from the responses of these two encoders under varying levels of response noise. To compare the encoders fairly, we normalized their outputs to have equal average power before adding noise. Consistent with our hypothesis, the quadratic encoder transmits more information about the song envelope than the threshold linear encoder (Fig. 5B, C). Thus, rather than being a by-product of quadratic encoding, frequency doubling enhances the representation of the song envelope.

### Quadratic filtering preserves a simple code for the song carrier

Quadratic encoding of the envelope raises an obvious question: can downstream neurons still recover the carrier? Linear filtering preserves stimulus frequencies, making carrier reconstruction straightforward. By contrast, quadratic filtering generates pairwise combinations of input frequencies, including the frequency doubling observed in the CAP. For spectrally complex sounds, these additional frequencies create ambiguity because it becomes impossible to determine which frequencies were present in the original stimulus. Courtship song largely avoids this ambiguity because it is spectrally simple. Over the duration of the quadratic filter, both pulse and sine song are effectively narrow-band, and their dominant frequencies fall within the phase-locking regime of JONs (Fig. 5D,F; compare Fig. 2H–K). In this regime, the quadratic filter’s main effect is that of frequency doubling—the dominant frequency in the response is twice the stimulus carrier frequency. Consequently, the carrier frequency can be recovered simply as half the dominant response frequency for both sine and pulse song [42] (Fig. 5E, G, S7). Consistent with this prediction, downstream auditory neurons exhibit sharp frequency tuning [32, 62, 63].

### Intensity adaptation for song enables robust pattern recognition

Having shown that quadratic filtering preserves information about both the song envelope and carrier, we next asked whether divisive normalization preserves these representations across the large intensity fluctuations encountered during natural courtship. This is relevant since sound intensity at the female changes drastically during courtship because of the angle and distance dependence of the arista [40, 64, 65]. To do this, we employed a classifier which uses the Euclidean distance between responses to classify the identity of short stimulus patterns across intensities [66]. We first quantified recognition of 100 short noise patterns, each presented at 8 different intensities (Fig. 6A). We analyzed both CAP responses and model responses, allowing us to isolate the contribution of divisive normalization by removing the DN stage (Fig. 6B,C). Responses from the CAP and the model with DN cluster primarily by stimulus identity rather than intensity (Fig. 6D, E, S8). By contrast, responses from the model lacking the DN cluster by intensity rather than pattern identity. Consequently, pattern information is high for both the CAP and model with DN, reaching ∼85 % of the theoretical maximum (Fig. 6F). For the model without DN, pattern information is strongly reduced. Thus, divisive normalization transforms an intensity-dependent neural representation into an intensity-invariant representation of stimulus identity. Courtship song poses a more demanding test because adaptation is incomplete: rapid intensity fluctuations can occur faster than the adaptation process itself [38]. We therefore classified stimulus identity for short song patterns of different intensities. The model with DN achieved near-perfect classification of song patterns, whereas removing DN markedly reduced performance (Fig. 6G, S8). Together, these results demonstrate that the strength and timescale of divisive normalization are well matched to the statistics of courtship song, enabling robust recognition of song patterns across the large intensity fluctuations encountered during natural social interactions.

**Figure 6:**
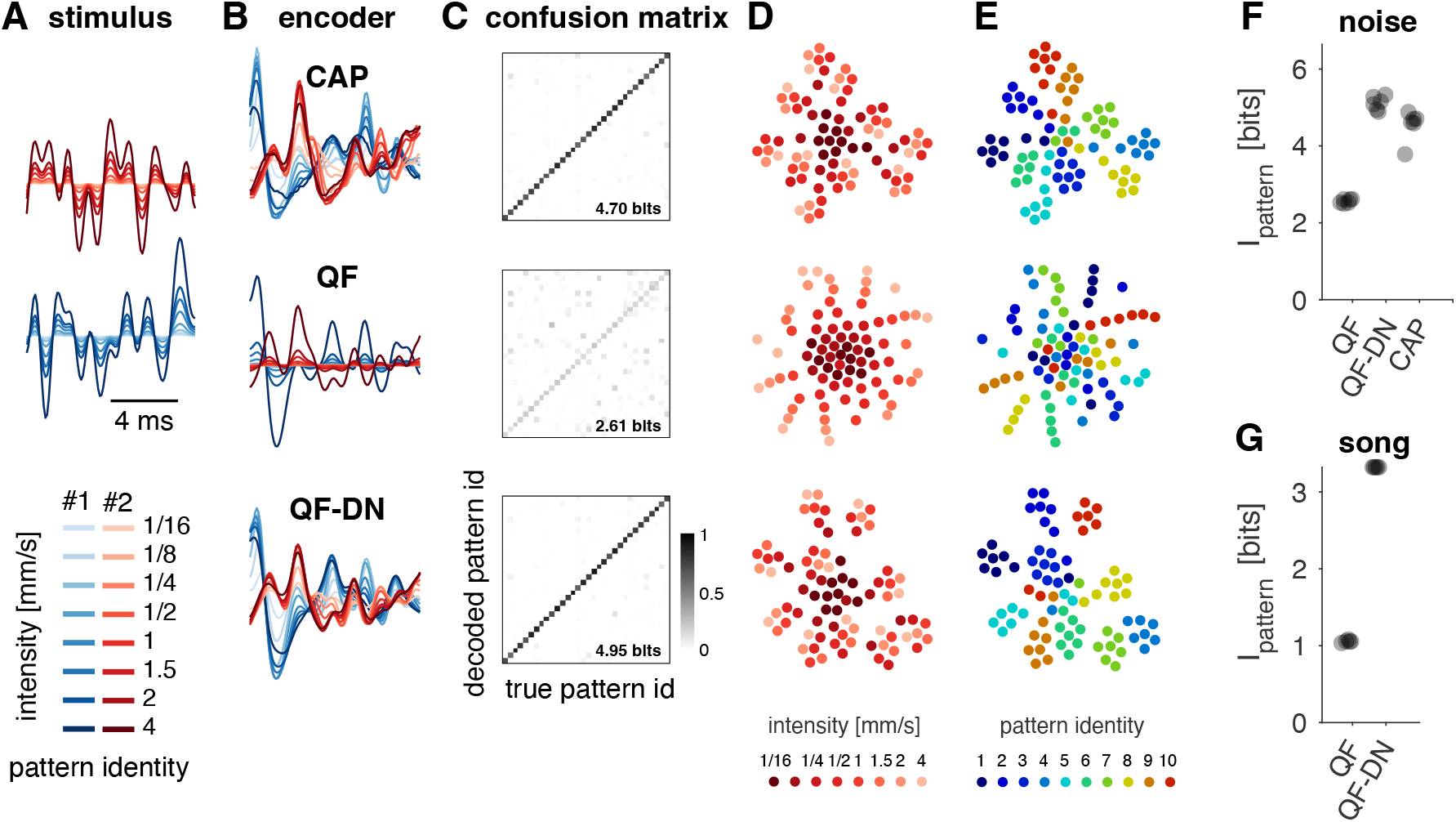
Divisive normalization improves pattern recognition across intensities. **A** Two different noise patterns (blue, red) at eight different intensities (color coded, see legend). **B** Responses to the noise patterns in A from the CAP (top) and from QF models with (QF-DN, bottom) and without (QF, middle) a divisive normalization stage. **C** Confusion matrices (color coded, see color bar) showing the classifier output for 4 of the 100, 10 ms-long noise patterns presented at 8 different intensities. Diagonal entries correspond to correct classification. Classification performance was quantified using mutual information in bits. **D, E** t-SNE visualization of the CAP, QF, and QF-DN responses for 10 different noise patterns, each presented at 8 different intensities. Responses are color coded by intensity (D) or pattern identity (E) (see legends). In the CAP and in a QF model with divisive normalization, responses cluster by pattern identity (E, top, bottom). In a model without divisive normalization, intensity is encoded by the stimulus magnitude, corresponding to the distance from the center of the point cloud (D, middle). **F, G** Pattern information across intensities for noise (F) and song (G) patterns. CAP and QF-DN responses contain more information about pattern across intensities than the QF responses. Dots correspond to individual flies (N=6).

## Discussion

Auditory receptor neurons in *Drosophila* implement multiple adaptive and nonlinear computations, including quadratic filtering, adaptive temporal filtering, and adaptation to stimulus mean and variance. We developed a unified computational model that accurately reproduces responses to both artificial and natural sounds using only two computational stages—a quadratic filter followed by divisive normalization (Figs. 1, 2). The model extends previous descriptions that focused on individual computations, such as active amplification [27, 48] or mean and variance adaptation [38], and provides a compact description of JON responses that is independent of the still unknown molecular implementation. Beyond reproducing JON responses, the model yields two central insights. First, it localizes distinct computations to different stages of auditory processing, separating adaptive antennal mechanics from neuronal gain control (Figs. 3, 4). Second, it reveals that computations which appear to generate an ambiguous and history-dependent code for arbitrary sounds instead produce a simple and robust representation of courtship song (Figs. 5, 6). Thus, the function of these computations can only be understood in the context of the statistics of behaviorally relevant signals. An important advantage of this computational framework is that it links electrophysiological measurements directly to computational building blocks. As a consequence, genetic or pharmacological perturbations can be interpreted in terms of changes in specific computations rather than simply changes in response amplitude or tuning.

### Mechanistic organization of adaptive and nonlinear processing

Our model clarifies how adaptive and nonlinear computations are distributed across the auditory periphery. Adaptive temporal filtering arises almost entirely from active antennal mechanics. Incorporating experimentally measured antennal filtering into the model eliminated the intensity dependence of the neuronal filter (Fig. 3), demonstrating that the increase in preferred frequency with sound intensity originates from adaptive mechanics rather than neuronal processes. This extends previous work using pure-tone stimulation [43, 47, 60, 61] by showing that adaptive antennal mechanics account for responses to arbitrary dynamic stimuli, including natural courtship song.

The remaining neuronal filter is largely intensity invariant (Fig. 3G), arguing against substantial contributions from spike generation, spike-frequency adaptation, or axo-axonal interactions to adaptive temporal filtering [67]. Instead, adaptive temporal filtering likely arises through regulation of mechanotransduction itself. One possibility is adaptation of the mechanotransducer channels, such as nompC. Alternatively, gain control may be mediated by accessory components of the transduction machinery, including inactive and nanchung, which are known to regulate the active mechanics of the antennal receiver [45, 68]. Recent work also implicates the voltage-gated potassium channel Shal (Kv4) in active antennal tuning and auditory mechanotransduction [69]. Although these possibilities remain speculative, the model considerably narrows the range of candidate mechanisms.

The model also constrains the origin of the quadratic nonlinearity that gives rise to frequency doubling (Fig. 2). Although frequency doubling has long been recognized in JON responses [37, 60], its cellular origin remains unresolved. Our analyses show that quadratic filtering acts downstream of adaptive mechanical filtering but upstream of divisive normalization (Fig. 3F,G).

Frequency doubling was originally proposed to arise from summation across two receptor populations with opposite directional preferences within Johnston’s organ [34]. In this model, one population responds to positive antennal displacements whereas the opposing population responds to negative displacements, together producing responses at twice the stimulus frequency. However, more recent anatomical and physiological evidence argues against this interpretation. Frequency doubling is observed even within subsets of JONs located on one side of Johnston’s organ [70], and adaptation transfers between positive and negative antennal displacements, suggesting that both response components arise within individual neurons [27].

Our model is consistent with this latter interpretation. Individual JONs may therefore respond to both directions of antennal displacement, either because mechanotransduction channels are coupled symmetrically to the cytoskeleton or because distinct channel populations respond preferentially to opposite displacement directions. Distinguishing between these possibilities will ultimately require recordings from identified individual JONs.

Our model further separates two mechanistically distinct components of variance adaptation. First, the delayed suppressive interaction within the quadratic filter suppresses sustained responses at high frequencies (Fig. 2H–L). This differentiating operation resembles the biphasic filters associated with spike generation in olfactory receptor neurons [12, 71] and could similarly arise from active conductances during spike initiation. Testing this hypothesis experimentally—for example by manipulating sodium-to-potassium conductance ratios—would directly examine whether spike generation contributes to the suppressive component of the filter.

Second, divisive normalization provides gain control across the entire frequency range (Fig. 4). Because inhibitory interactions between JONs are not known, divisive normalization likely arises within individual receptor neurons rather than through network interactions, in contrast to many other examples of divisive normalization [11, 72, 73]. Existing physiological data place variance adaptation downstream of mechanotransduction and partly after spike generation [38]. The short adaptation time constant measured here (Fig. 4C) suggests implementation by fast voltage- or calcium-dependent potassium conductances. Adaptation of subthreshold receptor potentials could arise through low-threshold potassium channels such as Shaker [74], whereas spike-frequency adaptation is more likely mediated by high-threshold channels including slowpoke [75] or ether-à-go-go [76]. More generally, fitting computational models to CAP responses from targeted genetic perturbations should provide a powerful strategy for assigning specific computations to individual molecular components.

### Adaptive and nonlinear computations simplify the representation of communication signals

Viewed individually, the computations implemented by JONs appear to produce a highly complex neural code. Quadratic filtering generates new frequency components, adaptive temporal filtering changes frequency tuning with stimulus intensity, and divisive normalization makes sound intensity coding history dependent. For arbitrary broadband sounds, these transformations indeed produce an ambiguous representation from which carrier frequency and envelope cannot easily be reconstructed.

Our analyses show that this apparent complexity largely disappears for the signals that flies actually use. Courtship song is spectrally narrow-band and occupies the frequency range in which the quadratic filter behaves as a frequency doubler rather than a canonical envelope detector (Fig. 2H–L). Consequently, carrier frequency can be recovered simply as half the dominant response frequency (Fig. 5D–G; [42]), while frequency doubling simultaneously improves representation of the song envelope by increasing its temporal sampling (Fig. 5A–C). Likewise, divisive normalization removes slow intensity fluctuations arising from changes in the relative position of the male and female during courtship [40, 64, 65] while preserving the faster envelope fluctuations that define pulse song (Fig. 6). Thus, computations that appear to increase ambiguity for arbitrary stimuli instead produce a representation of courtship song that is robust to noise, invariant to behaviorally irrelevant intensity fluctuations, and straightforward for downstream neurons to decode.

This organization also provides a unified explanation for the dual functions of hearing in flies. During acoustic threat detection, JONs emphasize rapid increases in sound energy through the combined action of quadratic filtering and adaptation, providing a simple code for downstream escape circuits [70]. During courtship, the same computations preserve both carrier and envelope information. Envelope information can be extracted by low-pass filtering, as observed in second-order auditory neurons [77–79], whereas carrier frequency remains explicitly represented through frequency doubling. The resulting code is therefore well matched to the natural statistics of communication signals [80].

### Efficient coding depends on behavioral goals

Our results illustrate a broader principle for sensory coding. Classical efficient coding theory emphasizes maximizing information transmission across ensembles of natural stimuli [4, 7–10, 81]. Judged by this criterion alone, JONs appear inefficient: quadratic filtering introduces ambiguity, adaptive filtering changes the representation with sound intensity, and divisive normalization discards information about absolute intensity.

However, animals rarely need to represent arbitrary natural stimuli. Instead, sensory systems evolved to solve specific behavioral tasks. In *Drosophila*, these tasks are dominated by acoustic communication and threat detection. Under these natural stimulus statistics, the computations implemented by JONs no longer reduce coding efficiency but instead emphasize the stimulus features most relevant for behavior while suppressing nuisance variables such as slow antennal movements and intensity fluctuations. In this sense, JONs are not optimized for reconstructing arbitrary sounds but for robustly extracting behaviorally relevant information.

More generally, our results suggest that the function of adaptive and nonlinear computations cannot be evaluated independently of the statistics of the signals they encode. Computations that appear unnecessarily complex—or even detrimental—when analyzed using generic stimulus ensembles become both simple and effective once considered in their ecological context.

## Methods

### Flies

Virgin females of the *Drosophila melanogaster* wild-type strain CantonS Tully were used for all recordings. Flies were sexed and housed in groups of ∼10 at 25 °C on a 12:12-h light-dark cycle. All experiments were performed 2–5 days post-eclosion.

### Stimulus design and presentation

Stimuli were generated at a sampling frequency of 10 kHz. Band-limited Gaussian noise (from now on termed “noise”) was produced from a sequence of normally distributed random values by band-pass filtering using a linear-phase, finite impulse response filter with a passband between 100 Hz and 1000 Hz. The effect of intensity on JON responses was estimated using 5 independent noise patterns that lasted 5 s and were presented at intensities of 1/16, 1/8, ¼, ½, 1, 1.5, 2, and 4 mm/s (“noise” stimuli in Fig. 1). For probing adaptation, we switched the intensity of the noise every 100 ms in a sequence that contained all transitions between ¼, ½, 1 and 2 mm/s (“step” stimuli in Fig. 1). The effect of adaptation on intensity tuning was assessed using a noise stimulus at intensities ¼, ½, 1, 2, 4 mm/s (adaptation background) whose intensity was switched every 120 ms for 20 ms to a probe intensity of 1/16, 1/8, ¼, ½, 1, 2, 4, and 8 mm/s (“background” stimuli in Fig. 1G, H). To minimize artifacts from abrupt changes in sound intensity, each intensity switch in the step and background stimuli had a duration of 1 ms during which the intensity was linearly interpolated to the new value. We also assessed responses to natural courtship, recorded from a *Drosophila melanogaster* male courting a virgin female [20, 21] (“song” stimuli).

### Sound

The sound delivery system consisted of i) the analog output of a DAQ card (PCI-6251, National Instruments), ii) a 2-channel amplifier (Crown D-75A), iii) a headphone speaker (KOSS, 16 Ohm impedance; sensitivity, 112 dB SPL/1 mW), and iv) a coupling tube (12 cm, diameter: 1 mm).

The stimulus presentation setup was calibrated as in Clemens et al. [78]. Briefly, the amplitude of pure tones of all frequencies used (100–1000 Hz) was calibrated using a frequency-specific attenuation value measured using a calibrated pressure gradient microphone (NR23159, Knowles Electronics Inc., Itasca, IL, USA). To ensure that the temporal pattern of the noise stimuli was reproduced faithfully, we filtered the presented noise patterns using the inverse of the system’s transfer function, measured using a pressure microphone (4190-L-001, Brüel & Kjaer) that was placed at the position of the fly’s head during recordings.

### Electrophysiology

Extracellular recordings were performed using glass electrodes (1.5/2.12 inner/outer diameter, WPI) pulled with a micropipette puller (Model P-1000, Sutter Instruments). The fly’s wings and legs were removed under cold anesthesia and the abdomen was subsequently fixed using low-melting-point wax. The head was fixed by extending and waxing the tip of the proboscis. The preparation was further stabilized by applying wax or small drops of UV-curable glue to the neck and the proboscis. The recording electrode was placed in the joint between the second and third antennal segments and the reference electrode was placed in the eye. Both electrodes were filled with external saline [82]. The sound delivery tube was placed orthogonal to the arista on the side from which we recorded JON activity, at a distance of 2 mm. The recorded signal was amplified and band-pass filtered between 5 Hz and 5000 Hz to reduce high-frequency noise and slow baseline fluctuations induced by spontaneous movement of the antenna (Model 440 Instrumentation Amplifier, Brownlee Precision). We ensured that the band-pass filter did not distort the recorded signal; e.g., it did not introduce artefactual response transients. We subsequently digitized the recorded signal at 10 kHz with the same DAQ card used for stimulus presentation (PCI-6251, National Instruments).

### Laser Doppler vibrometry

Arista movement was measured using laser Doppler vibrometry (Polytec OFV534 laser unit, OFV-5000 vibrometer controller, Physik Instrumente, low-pass 5 kHz).

## Data analysis

### Pre-processing

The instantaneous amplitude of the CAP was estimated from the envelope of the recorded signal as the magnitude of the Hilbert transform (Fig. S6A). Tuning curves from the “background” stimuli were obtained by averaging the CAP amplitude over the first 8 ms of the probe intensities (see stimulus description above).

### Adaptation time scale and strength

The adaptation time scale was estimated from the CAP amplitude traces by fitting an exponential function *r* (*t*) = *r*_*0*_ + *r*_max_ exp(*−t*/*τ*) to the falling/rising phases of positive/negative transients following intensity changes in the “step” stimuli (Fig. S6C).

## Modelling

### Model Structure

To reproduce the transfer function from stimulus waveform to CAP waveform, we used a discretized Volterra series truncated at second order, comprising a constant term *h*_*0*_, a linear term *h*_*1*_, a quadratic term *H*_*2*_, and a residual term *ϵ*(*t*):

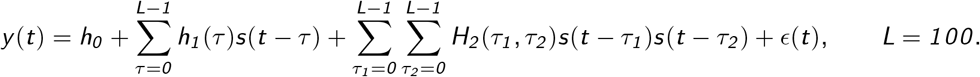

*s*(*t*) and *y* (*t*) are the stimulus and the CAP response, respectively. *h*_*0*_ describes a constant offset or bias. *h*_*1*_ is a linear filter and describes how stimulus values *τ* time steps into the past are weighted in the response. *H*_*2*_ is a quadratic filter and describes how products of stimulus values at two different time steps in the past, *τ*_*1*_ and *τ*_*2*_, are weighted in the response. The linear models in this paper only include the *h*_*0*_ and *h*_*1*_ terms. The quadratic filter model (QF) consists of all terms up to and including *H*_*2*_. The temporal support of the filters *h*_*1*_ and *H*_*2*_, which describes the memory of the system, was set to 10 ms (*L* = *100* samples). Model performance saturated at this duration— in initial tests, a longer temporal support did not improve model performance. This is confirmed by the lack of filter structure in *h*_*1*_ and *H*_*2*_ for *τ > 8* ms (Fig. 2A).

### Model fitting

The individual terms of the model—*h*_*0*_, *h*_*1*_, and *H*_*2*_ —were estimated using linear regression: *y*^*′*^(*t*) = ***σ***(*t*)**w**, where *y*^*′*^ is the predicted response and **w** is a concatenation of the model coefficients: [*h*_*0*_, **h**_*1*_, **h**_*2*_]. Exploiting the symmetry of the quadratic filter, *H*_*2*_ (*τ*_*1*_, *τ*_*2*_) = *H*_*2*_ (*τ*_*2*_, *τ*_*1*_), **h**_*2*_ is a vector containing all upper triangular values including the diagonal of *H*_*2*_ (*τ*_*1*_ *≥ τ*_*2*_). Equivalently, ***σ***(*t*) is a concatenation of the inputs for each of the model terms: [*s*_*0*_, **s**_*1*_, **s**_*2*_]. For the bias *h*_*0*_, the input *s*_*0*_ = *1*. For the linear filter *h*_*1*_, the input *s*_*1*_ (*τ*) corresponds to the stimulus in the 10 ms preceding the response time *t*: *s*_*1*_ (*τ*) = *s*(*t − τ*) for *τ* = *0*, …, *L − 1*. To reduce the number of filter coefficients to be estimated, we projected **s**_*1*_ onto a basis composed of 50 Gaussian bumps with a standard deviation of 0.2 ms (2 samples) and a spacing of 0.2 ms (2 samples), with the first bump at 0 ms and the last bump at 9.8 ms. For the input values to *H*_*2*_, we took the outer product of the stimulus values in the 10 ms (100 samples) preceding the response to be predicted: *S*_*2*_ (*τ*_*1*_, *τ*_*2*_) = *s*(*t − τ*_*1*_)*s*(*t − τ*_*2*_). To reduce the number of filter coefficients, we projected each matrix onto a basis composed of two-dimensional Gaussian bumps, each bump with a standard deviation of 0.3 ms (3 samples) and a spacing of 0.2 ms (2 samples) (cf. Rajan et al. [56]). We exploited the symmetry of *S*_*2*_ and kept only the unique values of *S*_*2*_ for which *τ*_*1*_ *≥ τ*_*2*_, flattened into a vector **s**_*2*_. The projection onto Gaussian bump bases and the exploitation of symmetry reduced the number of free parameters from *1* + *100* + *100 × 100* = *10, 101* to *1* + *50* + (*50*^*2*^ + *50*)/*2* = *1, 326*.

All filter coefficients were then fitted using ridge regression. Ridge regression minimizes the mean-squared error between the data and the predicted responses and the norm of the filter coefficients: 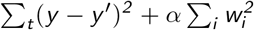 [83]. The first term increases the match between the data and the model, the second term penalizes filter weights that do not contribute to improving this match. *α* controls the influence of the penalty term and was chosen using methods and code from [84]. Since song is relatively sparse, we only used samples that were within *±100* ms of song for model fitting and evaluation. For visualization and analysis of the filters, *h*_*1*_ and *H*_*2*_ were projected from the basis of Gaussian bumps back into the temporal domain.

For linear-nonlinear models, which only contain the first two terms (*h*_*0*_, *h*_*1*_), we estimated an output nonlinearity based on methods described in [49]. For the quadratic models, the nonlinearity was approximately linear and did not improve performance. We therefore omitted that step to simplify model interpretation.

### Fitting the antennal filters and the antQF model

The transfer function from stimulus to antennal movement for each intensity was fitted as a discretized Volterra series with terms *h*_*0*_ and *h*_*1*_. This model typically explained more than 90 % of the variance in antennal responses to noise (Fig. S4C). To account for intensity-dependent antennal filters in the quadratic model, the intensity-specific antennal filters obtained from one animal were used to pre-filter the stimulus before fitting the quadratic model. This was necessary because simultaneous measurement of antennal movement and CAP responses was not possible in our rig. However, antennal filters are virtually identical across animals (pairwise Pearson correlation of the filters *r* = *0* .*97 ± 0* .*03* across *N* = *5* flies; filters from different intensities were concatenated before calculating the correlations).

### Fitting QF-DN

The divisive normalization (DN) stage has the form *x*^*′*^ = *x* /(*1* /*θ* + *γ*), where *γ* is the gain-control signal that divides the input *x*and is given by 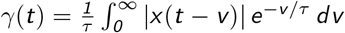. *τ* is the adaptation time constant. The adaptation kernel was normalized to have an integral of 1.0 to make the magnitude of the gain control signal independent of *τ*. The input to the DN stage, *x*, is the output of the quadratic filter. The QF-DN model was fitted using an iterative procedure: We initialized the filter by fitting a QF model without the DN stage to the data using ridge regression. Then, we optimized the parameters of the DN stage, *θ* and *τ*, by minimizing the mean-squared error between the CAP and the model prediction using the MATLAB function “fmincon”, holding the filter coefficients constant. Lastly, we optimized the filter coefficients, holding the DN parameters constant. The model parameters typically converged after 1–2 cycles of fitting the parameters of the DN stage and the filter. During fitting, only the magnitude, not the structure, of the filter changed from its initialization.

### Estimation of nonlinearities

The nonlinearity for the LN model shown in Fig. 1C was calculated by filtering the stimulus with the linear filter. We then binned the filtered stimulus values, choosing the 24 bin edges such that each bin was populated with the same number of samples. The x and y values of the nonlinearity were given by the average filtered stimulus and CAP values for each bin.

### Model evaluation

Model performance was quantified using the squared Pearson correlation coefficient between the CAP and the model predictions. Initial experiments with cross-validation showed that training and test performance are within *±1* % of each other.

To further rule out overfitting, we compared model performance using the Bayesian Information Criterion, which penalizes models with more parameters: BIC = *N* ln(*E*) + *K* ln(*N*) [85]. E is the mean-squared error between the CAP and the model predictions, K is the number of model parameters plus 1 for the error variance, and N is the number of samples.

### QN representations

To gain insight into the computational structure of the quadratic filter, we represented *H*_*2*_ using its eigenvalue decomposition: 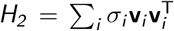, where the **v**_*i*_ and *σ*_*i*_ are the eigenvectors and their associated eigenvalues [51, 52]. This representation is useful, since *H*_*2*_ is typically low rank; that is, a few eigenvector-eigenvalue pairs are sufficient to reconstruct *H*_*2*_ with sufficient fidelity (Fig. S1). This representation is equivalent to a bank of linear-nonlinear models with filters **v**_*i*_ and quadratic nonlinearities 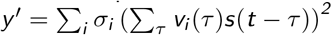.

### Filter cutoff frequency

The cutoff frequency of linear filters was computed in the frequency domain by first computing the filter’s power spectrum and then determining the frequency at which the power spectrum falls below 0.5 of its maximum.

### Subspace overlap

To compare quadratic filters fitted to different stimuli in a manner that is robust to noise, we computed the overlap between the subspaces spanned by the four leading eigenvectors. Specifically, given two quadratic filters *A* and *B*, with eigenvectors 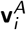 and 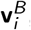, the cumulative overlap between the pairs of the *K* largest eigenvectors is given by 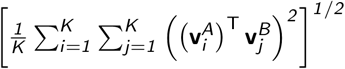 (Fig. S3D) [86].

### Envelope reconstruction

To assess the contribution of the quadratic filter to encoding the envelope, we compared the performance of two simple encoders: a rectified-linear encoder, which thresholds the stimulus at 0, and a quadratic encoder, which squares the stimulus. For a stimulus that is symmetric around 0, the quadratic encoder produces more response energy because the rectified-linear encoder cuts off all negative stimulus components. We therefore normalized the output of each of the encoders to unit norm. Gaussian noise with standard deviation *σ* was then added to the normalized outputs and the coherence between the model output, *r*, and the original envelope, *s*, was estimated from the stimulus: *C*_*sr*_ (*f*) = *P*_*sr*_ *P*_*rs*_ /(*P*_*ss*_ *P*_*rr*_) for different noise-to-signal ratios as a measure of decoding accuracy.

### Pattern classification

For pattern classification, we used a leave-one-out nearest-neighbor classifier. Each of the 100, 10 ms-long response patterns was selected as a template and assigned to the class of its nearest neighbor, based on the Euclidean distance to the 99 non-template patterns [87]. For the song, we used 10 different response patterns, each lasting 100 ms. The longer patterns were chosen because 10 ms barely exceeds the duration of a single song pulse. The classification resulted in a confusion matrix *p*(*s, s*^*′*^), which tabulates the joint probability with which a response elicited by stimulus *s* was classified as having been elicited by one of the stimuli *s*^*′*^. The mutual information of *p* serves as a lower bound on the mutual information between stimulus and response: 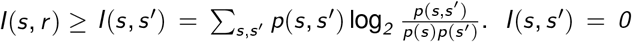 if and only if *p*(*s, s*^*′*^) = *p*(*s*)*p*(*s*^*′*^); that is, if the actual and classified stimulus identities are statistically independent. The maximal value of *I*(*s, s*^*′*^) is given by the log_*2*_ of the number of stimulus classes (log_*2*_ (*100*) = *6* .*64* for noise patterns, log_*2*_ (*10*) = *3* .*32* for song patterns, log_*2*_ (*8*) = *3* for intensity) for a perfect, one-to-one mapping between actual and classified stimulus. We visualized the similarity structure of the representations using t-SNE of a random subset of 10 patterns [88].

## Acknowledgements

We thank Cyrille Girardin for discussions during an early phase of the project; Georgia Guan for technical assistance; members of the Murthy lab, the Clemens lab, and Tim Gollisch for helpful feedback; and Rachel Wilson and Stephen Holtz for feedback on the manuscript. JC was supported by a postdoctoral fellowship through the Princeton Sloan-Swartz Center; an Emmy Noether grant (329518246) from the DFG; and DFG SFB889, Mechanisms of Sensory Processing (grant #154113120, Project A10). MM was supported by an NIH New Innovator Award (DP2) and a HHMI Faculty Scholar award.

## Author Contributions

Conceptualization: JC, MM.

Data collection and curation: JC.

Formal analysis: JC.

Funding acquisition: JC, MM.

Investigation: JC.

Methodology: JC, MM.

Project administration: MM.

Resources: MM.

Software: JC.

Supervision: MM.

Visualization: JC.

Writing—original draft: JC.

Writing—review & editing: JC, MM.

## Data Availability Statement

The data and code were deposited at https://doi.org/10.5281/zenodo.22173263 (https://github.com/janclemenslab/quadratic-adaptive).

## Ethics statement

Ethical approval was not required for this study because all experiments were conducted using *Drosophila melanogaster*, an invertebrate species not covered by the U.S. Public Health Service Policy on Humane Care and Use of Laboratory Animals or the U.S. Animal Welfare Act. All procedures were conducted in accordance with applicable institutional guidelines.

## Supplementary Information

**Figure S1:**
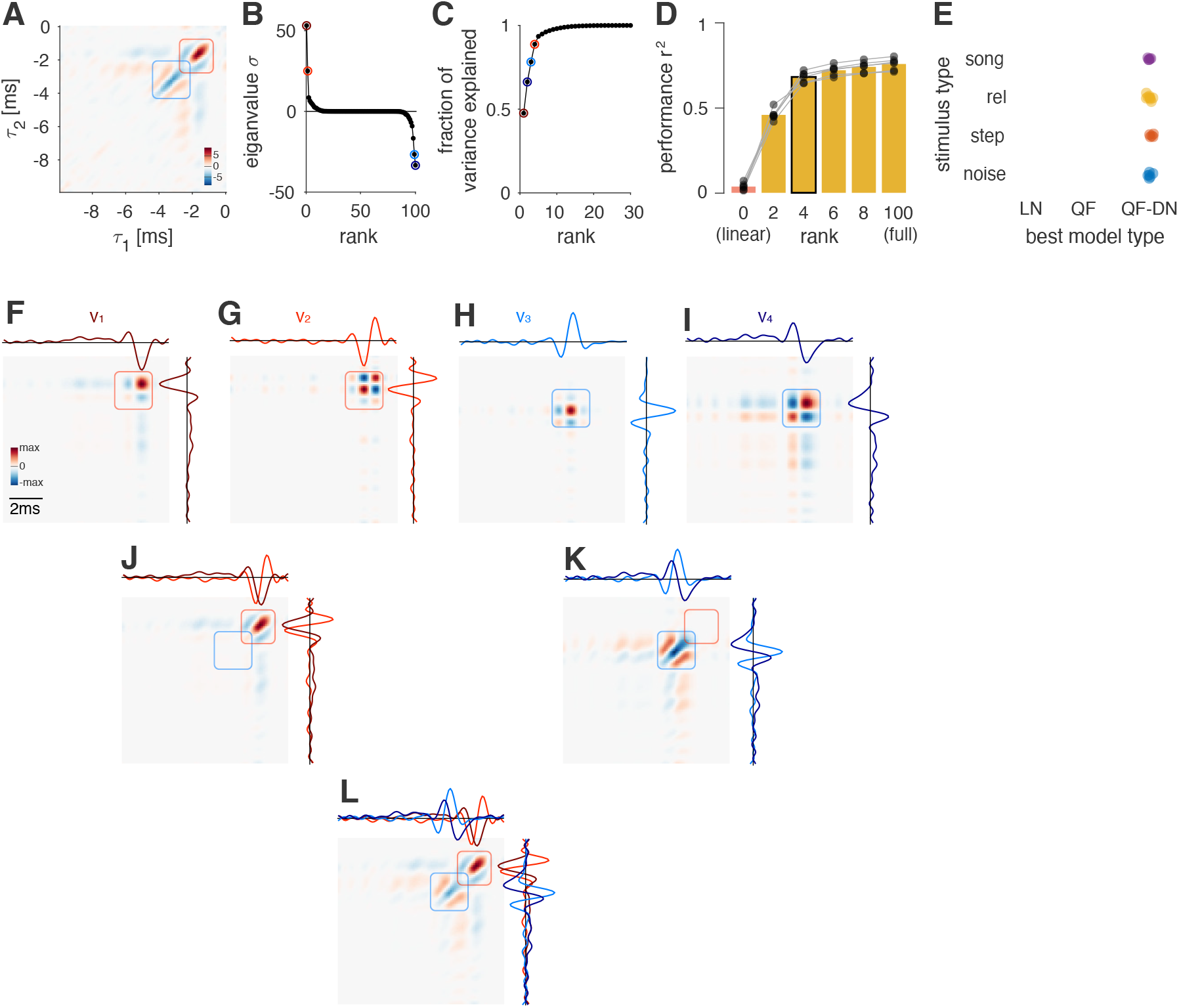
Eigenvalue decomposition and low-rank approximation of the quadratic filter *H*_*2*_. **A** Quadratic filter fitted to a noise stimulus (intensity 0.5 mm/s). **B, C** Eigenvalue spectrum of *H*_*2*_ (B) and cumulative sum of squared eigenvalues (C). The latter corresponds to the fraction of variance in *H*_*2*_ explained. The four eigenvalues with the largest absolute values are marked in color. **D** Performance of QF models using different low-rank approximations of *H*_*2*_. The rank corresponds to the number of eigenvalues used for representing the quadratic filter, equivalent to the number of LN models in the filter bank. The leftmost bar (pink) corresponds to a rank-0 model that contains only a bias and linear term, but no quadratic terms. A rank-4 model (with 4 LN models, black outline) is the simplest model with performance close to that of the original full-rank model. Bars show mean over flies and dots show values for individual flies connected by lines (N=6 flies). **E** Model with the lowest score of the Bayesian Information Criterion (BIC) for the four stimulus types. The BIC score is a measure of performance that penalizes models with more parameters. The QF-DN model has the lowest BIC score for all stimuli, which confirms the result of our *r* ^*2*^ -based performance analysis (Fig. 1G, H). Individual points correspond to flies. Numbers of flies for noise/step/background/song are 6/5/9/5. **F–I** Outer product matrix, 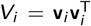, of each of the four leading eigenvectors. The waveforms aligned with the x and y axes show the eigenvectors **v**_*i*_ (right). **J, K** Sum of the outer-product matrices (weighted by the eigenvalues) for the excitatory (J, *σ*_*1*_ *V*_*1*_ + *σ*_*2*_ *V*_*2*_) and suppressive (K, *σ*_*3*_ *V*_*3*_ + *σ*_*4*_ *V*_*4*_) eigenvector pairs. The matrix in K is sign-inverted, since *σ*_*3*_ and *σ*_*4*_ are negative (see B). **L** A rank-4 approximation, given by the sum of all 4 outer products 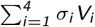, yields a good reconstruction of *H*_*2*_ (compare

**Figure S2:**
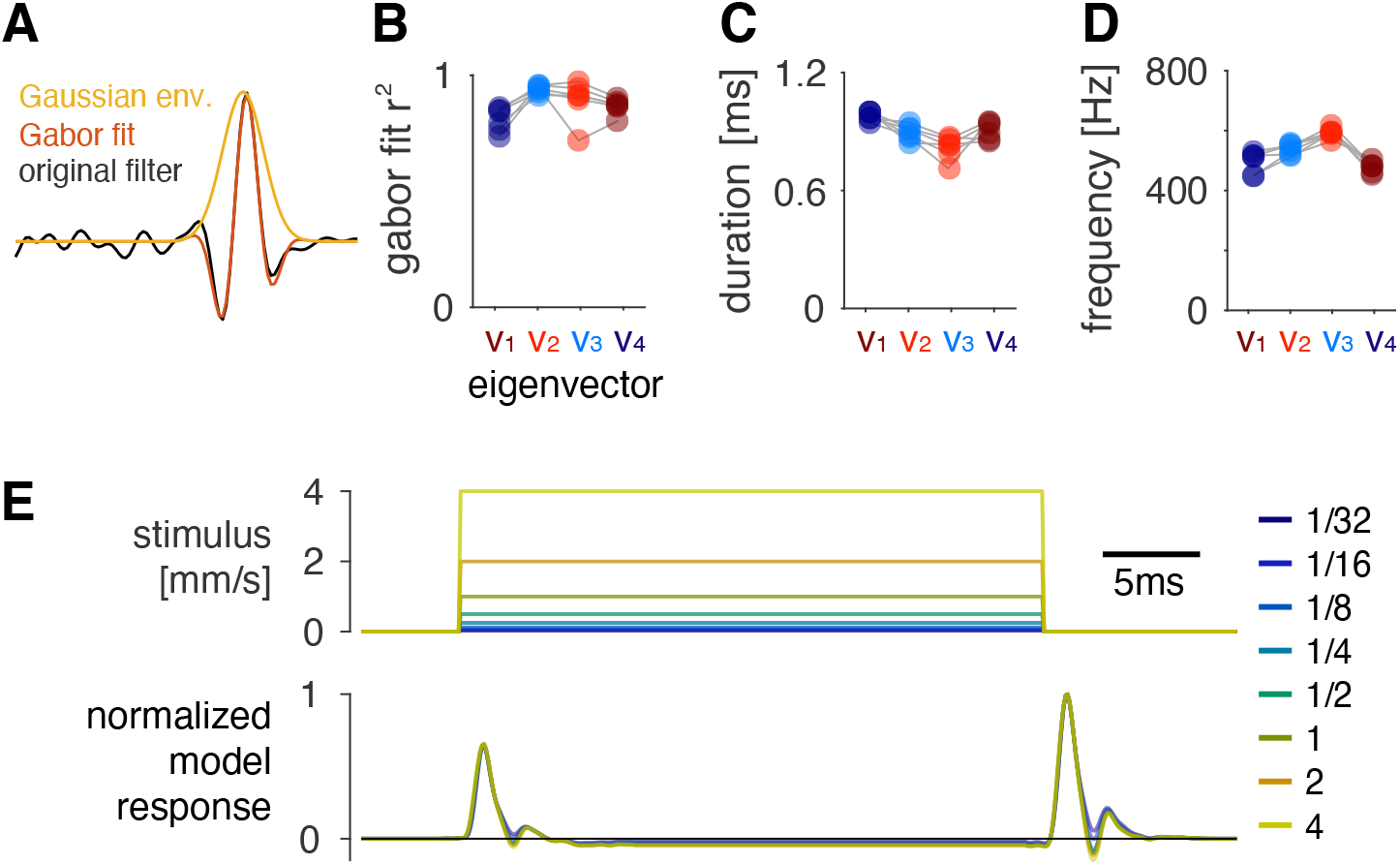
Performance, Gabor parameters, and responses to static stimuli. **A** The waveforms show an example fit (red) to one of the filters (black). The orange trace shows the Gaussian envelope of the filter. **B** The performance *r* ^*2*^ for the fits of the Gabor filters to the eigenvectors **v**_*1*_ –**v**_*4*_ of *H*_*2*_. **C, D** Duration (C) and carrier frequency (D) of the Gabor filters fitted to **v**_*1*_ –**v**_*4*_. Dots show recordings of individual flies, connected by lines (N=6 flies) in B–D. **E** QF output (bottom, normalized to peak at 1.0) for the static step stimuli with different intensities (top, intensity is color coded, see legend). The QF responds to the onsets and offsets of the step stimuli. This lack of responses to static stimuli is a correlate of mean adaptation.

**Figure S3:**
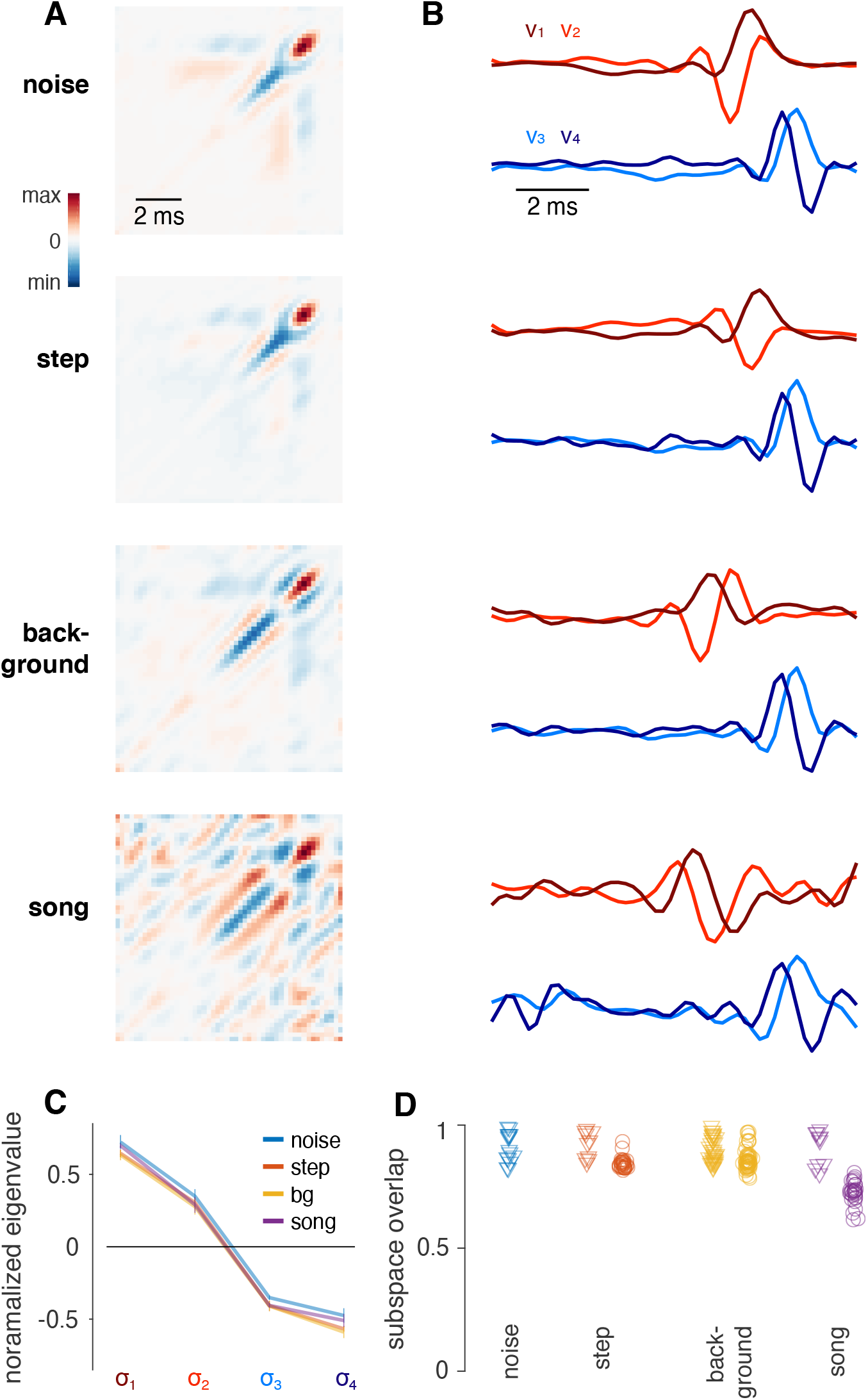
The filters of models fitted to different stimuli are similar. **A, B** Quadratic filter (A) and four leading eigenvectors **v**_*1*_ –**v**_*4*_ (B) for the four stimulus types used in this study. **C** Normalized eigenvalues of the QFs in A for the four stimulus types (“bg” = “background” stimulus). The four eigenvalues for each stimulus type were normalized to unit norm to account for differences in filter magnitude. Lines and error bars indicate mean ± std over flies. The relative amplitude of the eigenvalues is independent of stimulus type. **D** The subspace overlap measures the match of the hyperplane spanned by the four filters of models fitted to different stimuli (B). We take the filters fitted to noise as the reference. Triangles correspond to the subspace overlap for filters estimated for the same stimuli in different flies and reflect inter-individual variability. Circles indicate the overlap of filters estimated for different stimuli with those estimated for noise. N=6/5/9/5 flies for noise/step/background/song. The filter structure is robust to the stimulus type, indicated by the strong overlap between the subspaces spanned by filters fitted to different stimuli.

**Figure S4:**
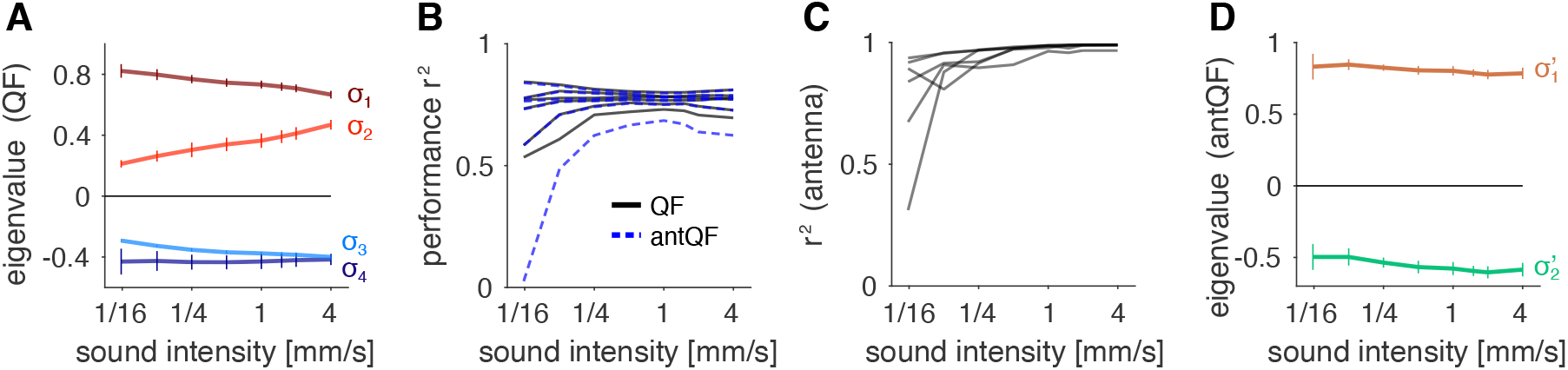
Model structure and performance for different intensities. **A** Eigenvalues of the QF, *σ*_*1*_ –*σ*_*4*_, fitted to noise stimuli of different intensities. **B** Performance (*r* ^*2*^) of the QF (black) and the antQF (blue) for different intensities. Each line marks the performance for a given fly. Except for one outlier recording both models perform identically, highlighting the computational equivalence of the QF and the antQF models. **C** Performance of the linear filter model that describes the transformation from sound stimulus to antennal position for each intensity. Each line marks the model performance for a given fly across intensities. **D** Eigenvalues *σ*^*′*^_*1*_ (brown) and *σ*^*′*^_*2*_ (green) of the antQF model for different intensities. Lines and error bars in A and D indicate mean ± SD over 6 flies.

**Figure S5:**
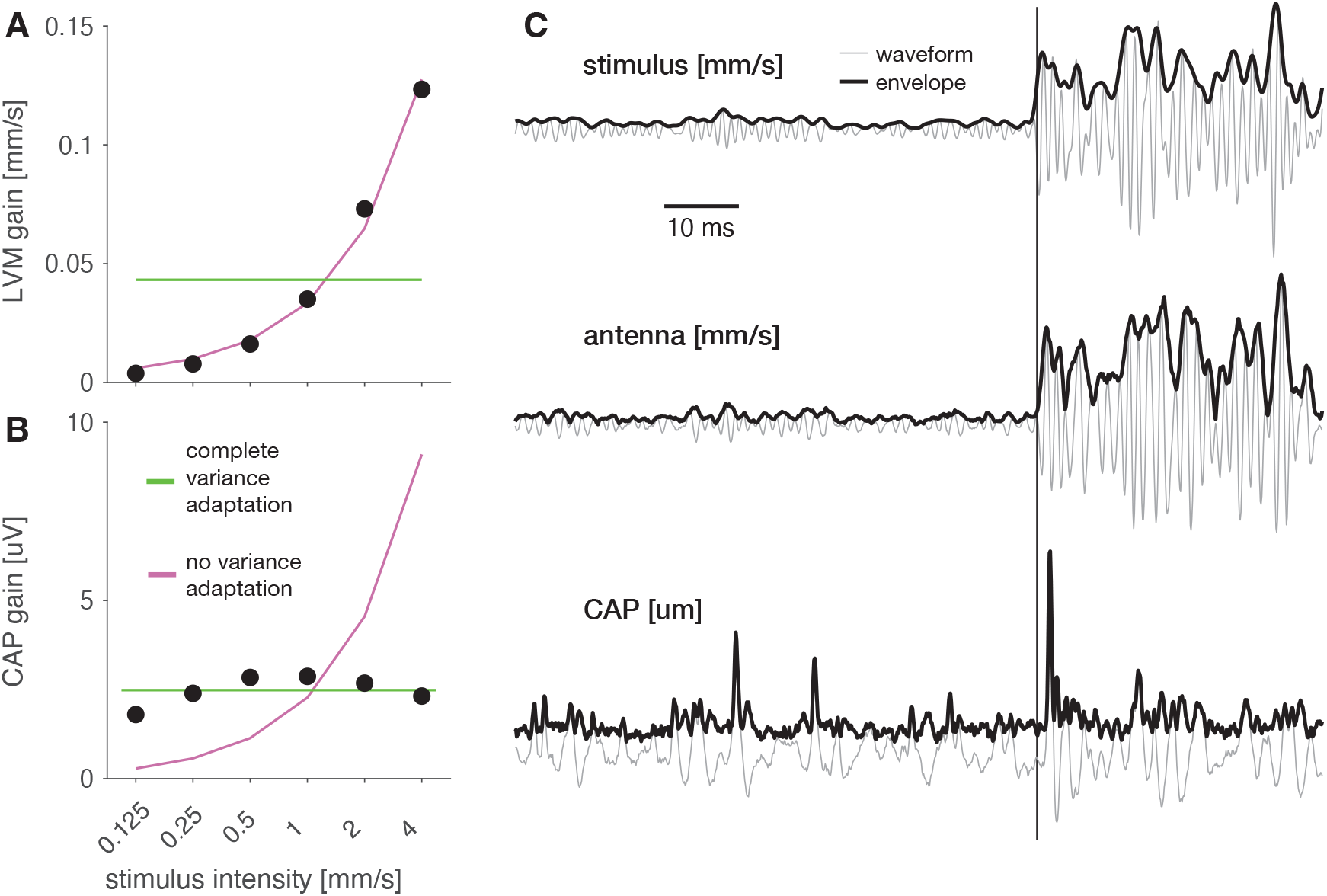
Antennal mechanics do not contribute to variance adaptation. **A, B** The gain (standard deviation) of responses from the antenna (A) and the CAP (B) to noise stimuli with different intensities. While CAP responses change little with intensity, indicating intensity invariance (horizontal green line) created by variance adaptation, antennal responses linearly scale with the intensity (purple line), indicating a lack of variance adaptation. **C** Responses of the antenna (middle) and the CAP (bottom) to a step in noise intensity (top). Thin grey lines indicate the signal waveform and thick black lines the signal envelope. Only the CAP produces transient responses to the intensity step, indicating a lack of variance adaptation in the antenna.

**Figure S6:**
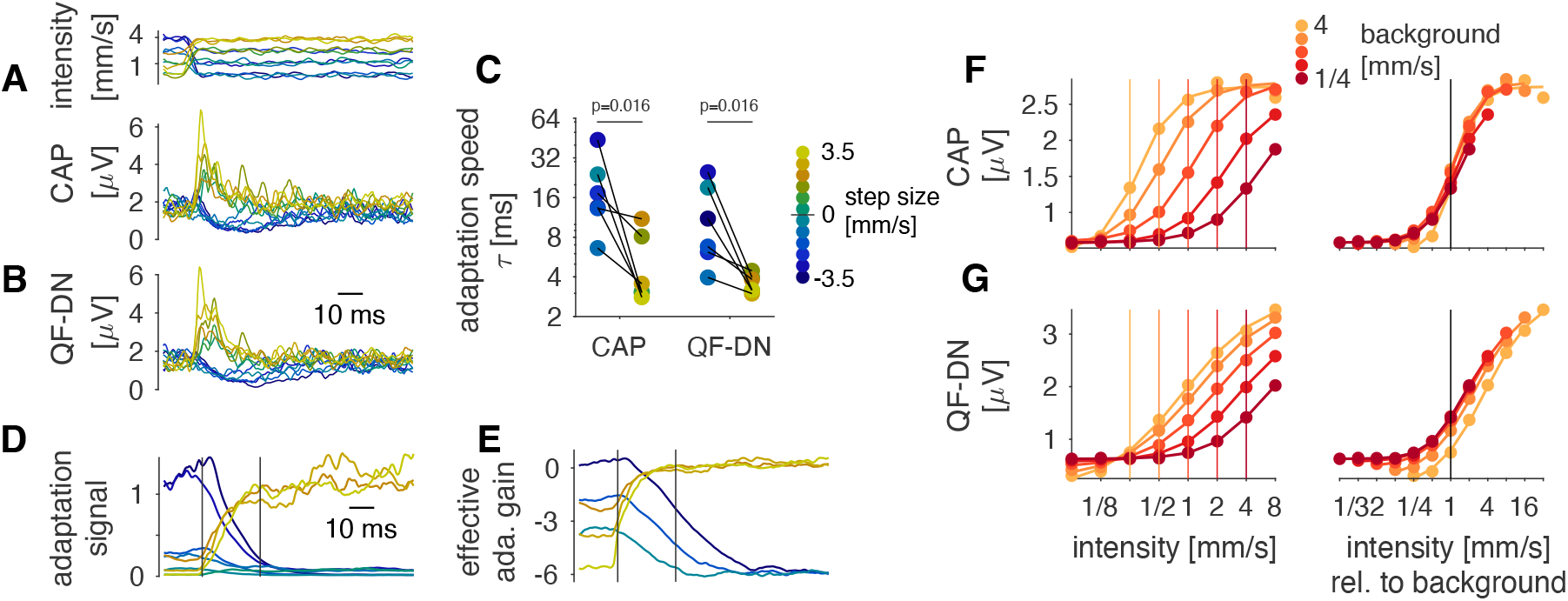
The model reproduces the asymmetrical dynamics and the sensitivity effects of adaptation. **A–B** Response magnitudes (CAP envelopes) for white noise with intensity steps (top) in the CAP (A) and the QF-DN model (B). Both the CAP and the model exhibit asymmetrical adaptation dynamics: Relaxation to steady state is fast for positive steps (yellow hues) and slower for negative steps (blue hues). See legend in C for color code. **C** The effective adaptation time constants in CAP and model responses are larger—adaptation is slower—for negative steps than for positive steps (blue and yellow points, respectively; see legend). Note that this is despite the model having a single, fixed parameter for adaptation speed (B). P-values from one-sided sign test (matched pairs are stimuli with same step size with different signs, black lines) for model and data. **D, E** Linear adaptation signal (D) and the effective adaptation gain (E), given by the log of the adaptation signal for the model responses in B. Vertical lines in D mark the beginning and end of the adaptation transients from the model in D. While the adaptation signal itself is relatively symmetrical for increases and decreases in intensity (D), the effective adaptation gain—proportional to the logarithm of the adaptation signal—exhibits the asymmetrical dynamics of the CAP (E). **F, G** Intensity tuning curves of the CAP (F) and the QF-DN model (G), adapted to different background intensities (color code, see legend). Tuning curves are plotted on an absolute intensity scale (left, vertical colored lines indicate background intensities) and with intensity given relative to the background (vertical black line at 1.0 depicts the background). In both the model and the data, tuning curves shift in proportion to the background intensity.

**Figure S7:**
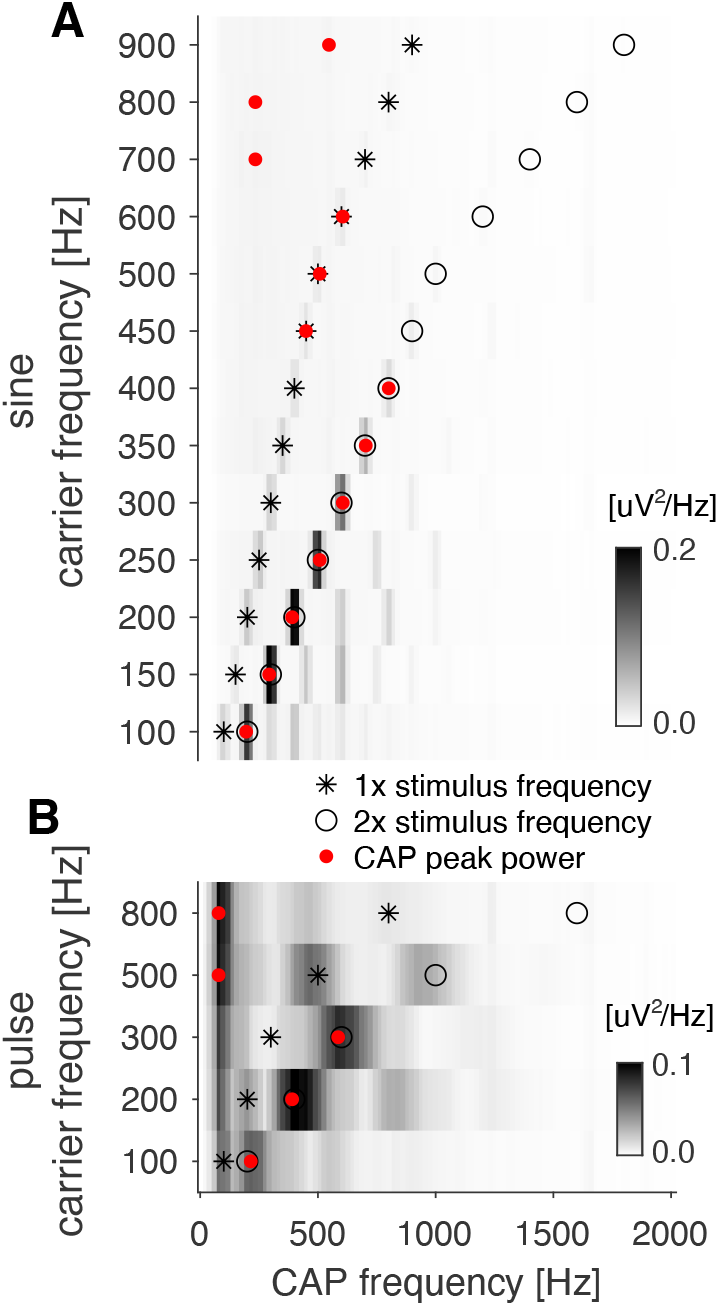
Decoding the carrier frequency of sine and pulse song from CAP responses. **A, B** CAP power spectra for sine tones (A) and pulses (B) with different carrier frequencies (y-axis) (spectral power of the CAP is color coded, see color bars). The carrier frequency and its double are marked as stars and open circles, respectively. The frequency of maximum CAP power is marked with a red dot. See Fig. 5D–G. In the frequency range occupied by song (100–500 Hz), the CAP oscillates at twice the carrier frequency for both pulse and sine. For higher frequencies, responses are either weak (A) or oscillate at very low frequencies (B).

**Figure S8:**
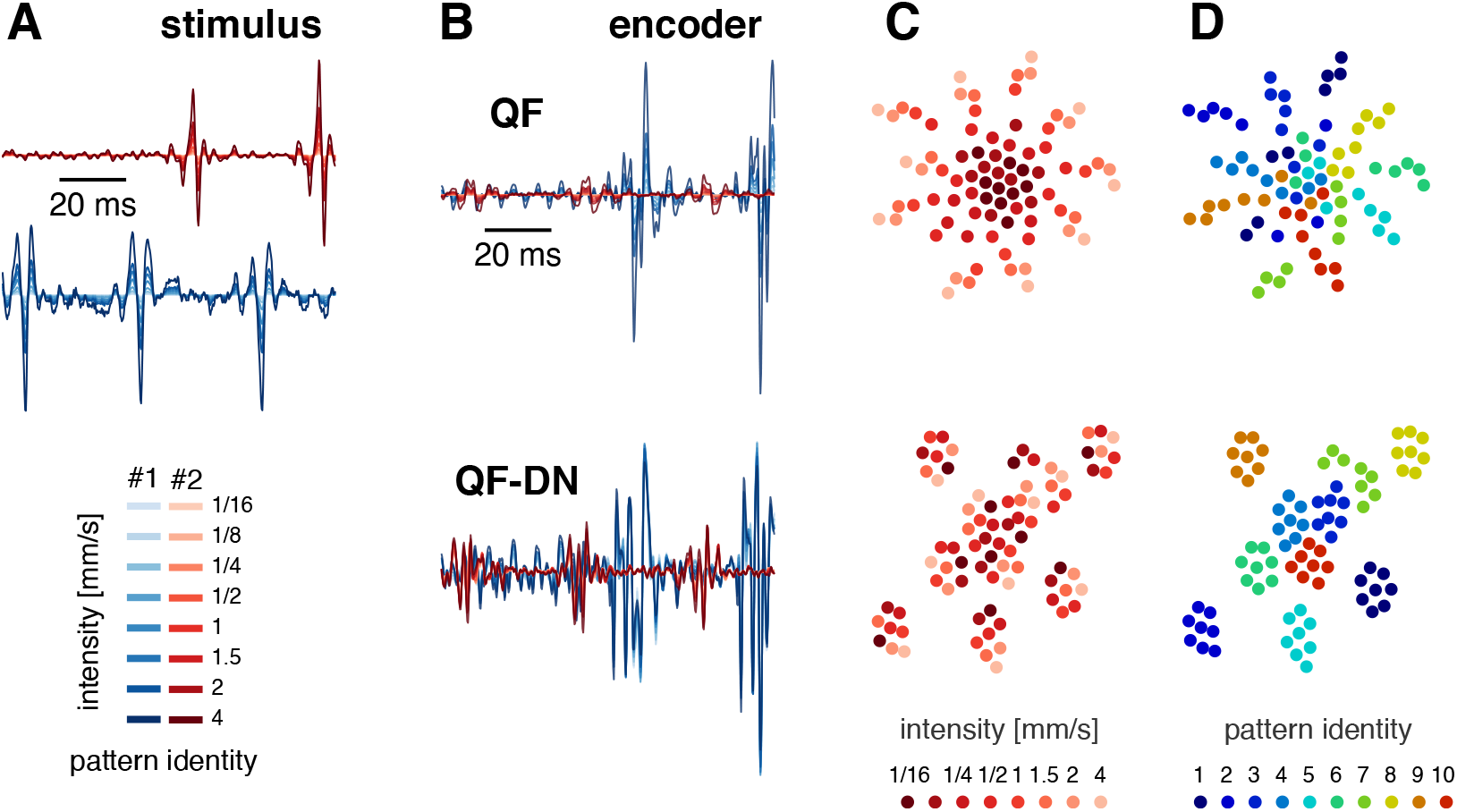
Divisive normalization improves pattern recognition across intensities for song. **A** Two different song patterns (blue, red) at eight different intensities (color coded, see legend). **B** Responses to the song patterns in A from QF models with (QF-DN, bottom) and without (QF, top) a divisive normalization stage. **C, D** t-SNE visualization of the QF and QF-DN responses for 10 different song patterns, each presented at 8 different intensities. Responses are color coded by intensity (C) or pattern identity (D) (see legends). In the QF model with divisive normalization, responses cluster by pattern identity (D, bottom). In the model without divisive normalization, intensity is encoded by the stimulus magnitude, corresponding to the distance from the center of the point cloud (C, top).

